# Ecological divergence and post-eclosion brain development shape visual performance during *Heliconius* speciation

**DOI:** 10.64898/2026.04.14.718491

**Authors:** José Borrero, Amaia Alcalde-Anton, Leo Laborieux, Daniel Shane Wright, Daniela Lozano-Urrego, Geraldine Rueda-Muñoz, Carolina Pardo-Diaz, Camilo Salazar, Stephen H. Montgomery, Richard M. Merrill

## Abstract

Sensory adaptation is increasingly recognized as a key driver of ecological speciation, but how visual system divergence is coordinated across development, and how this translates into behavioral differences, remains poorly understood. The butterfly *Heliconius cydno*, which inhabits closed-canopy forests, has larger eyes and greater investment in visual brain centers than its sympatric close-relative *H. melpomene*, which occupies more open forest-edge habitats, suggesting divergent ecological selection on the visual system. However, the behavioral consequences of these visual adaptations, their developmental trajectories, and whether they break down in hybrids is unknown. To address these questions, we combined ecological field data with behavioral assays and deep-learning-assisted segmentation of neuroanatomy. Visual acuity – the ability to resolve spatial detail – was higher in *H. cydno*, consistent with its greater ommatidia number, but also increased with age in both species despite no change in external eye morphology. These improvements coincided with the onset of male courtship and female oviposition, suggesting that early adult neurodevelopment shapes visual performance and may support the demands of reproduction. Brain morphology showed species-specific trajectories of post-eclosion optic lobe growth that broadly paralleled increases in acuity and were accompanied by ongoing neurogenesis in the adult optic lobes. While hybrids exhibited intermediate visual acuity, relationships among different components of the visual system were disrupted in hybrids. Together, these results show that ommatidia number alone cannot explain variation in visual acuity, and highlight how coordinated sensory evolution, and its breakdown in hybrids, may contribute to divergence during the early stages of speciation.

**Significance Statement:** Adaptation of sensory systems is increasingly recognized as a key driver of species formation, but it is unclear how these systems develop and shape behavioral differences. Using closely related *Heliconius* butterflies adapted to different light environments, we show that visual acuity improves during early adulthood through neural development, despite no change in eye structure. This maturation coincides with the onset of reproductive behaviors and is accompanied by growth and neurogenesis in visual brain regions. In hybrids, coordination among sensory traits breaks down, resulting in intermediate visual performance. These results show that sensory development is critical for behavioral adaptation and suggest that its integration contributes to the maintenance of species boundaries.

## Introduction

When the reliability and salience of sensory cues vary across environments, natural selection is expected to favor divergence in the sensory systems through which they are perceived (1–3). This may contribute to reproductive isolation between species: Pre-mating barriers can arise through selection against poorly adapted immigrants, or by influencing mate choice; and post-mating barriers may emerge if hybrids with intermediate sensory phenotypes suffer fitness costs (4). The best-known examples include rapid divergence in visual sensitivities in cichlid fishes (5), but adaptive divergence is now broadly documented across sensory systems, from changes in the sensory periphery to shifts in brain composition, in both aquatic and terrestrial taxa (6–10). However, we still know little about how selection in response to contrasting sensory environments is coordinated across different components of the sensory systems, or how this translates into behavioral differences across individual lifetimes.

Sensory systems comprise multiple interacting components, from peripheral receptor cells that capture environmental stimuli, to central neural circuits that process and integrate this information (11). To function effectively, sensory adaptations depend on these elements staying coordinated throughout development and evolution (12, 13). Rather than reflecting strict developmental constraints, this coordination can arise when selection acts on a functionally integrated system (14, 15). As such, if diverging populations accumulate heritable differences in these coordinated systems, hybridization may disrupt their integration (16, 17), leading to mismatches and potentially impairing behavioral performance (4). Determining whether ecological divergence generates integrated sensory architectures, and whether these break down in hybrids, is important for our understanding of how sensory ecology can influence species divergence.

In particular, although the eye and brain are functionally linked, their developmental and evolutionary trajectories can diverge, allowing selection to act on neural variation and shape visual performance independently of eye morphology (14, 18). Neural development and remodeling can extend into adulthood across vertebrates and invertebrates, with age- and experience-dependent changes reflecting ongoing refinement of sensory and cognitive function (19–21). In insects, this is best documented in the mushroom bodies, brain regions involved in associative learning and memory (18, 22–25). However, age-related development also occurs in the optic lobes, the primary visual processing centers, across a range of insects (14, 23, 24, 26–29), but its consequences for visual performance are not well characterized.

*Heliconius* butterflies are well known for their diversity of bright warning patterns, though speciation within the genus is often associated with broader ecological and behavioral shifts (30, 31). Vision plays a central role in many of these behaviors, including mate recognition, foraging and host plant selection (32–36). While extensive research on spectral sensitivity and color vision has revealed variation in photoreceptor composition across the genus (37–42), other aspects of visual perception remain less well explored. In particular, visual acuity, the ability to resolve fine spatial detail, is comparatively understudied (43). To date, only a single study has behaviorally quantified visual acuity in *Heliconius* (44), leaving unresolved how this aspect of visual performance varies between species inhabiting contrasting sensory environments, or across development.

The closely related species *Heliconius cydno* and *H. melpomene* provide an excellent system to study sensory adaptation. Although broadly sympatric in Central and northern South America, and ongoing gene-flow due to –albeit very rare– hybridization (45), they are ecologically segregated, with species boundaries additionally maintained by strong assortative mating, which involves a strong visual component (34, 36, 46, 47). *H. cydno* occupies closed-canopy forest, whereas *H. melpomene* is found in more open forest-edge habitats (48–50), environments that differ in light intensity, spectral composition, and visual complexity (49, 51). Despite similar spectral sensitivities based on opsin expression (38), these species show heritable divergence in eye morphology and neuroanatomy, with *H. cydno* possessing larger eyes and greater relative investment in visual brain regions than *H. melpomene* (6, 8). Such differences are likely functionally significant, as visual acuity is predicted to scale with ommatidia number (52). However, the behavioral consequences of this divergence, particularly for visual acuity, and how these traits develop across ontogeny, have not been tested. In *Heliconius*, substantial post-eclosion brain growth has been documented (53), alongside proliferative activity in the adult optic lobes (23). Nevertheless, it remains unclear whether this early adult development in the optic lobes contributes to the maturation of visual performance, or whether these species follow distinct developmental trajectories shaped by their contrasting ecological demands.

Here, we use an integrative approach to examine how ecological divergence and post-eclosion development shape visual performance in *H. cydno cydno* and *H. melpomene martinae*. We first confirm that at our field site in Colombia these species occupy distinct forest microhabitats associated with contrasting sensory environments. We then ask: (i) Does visual acuity differ between species, and is this variation linked to differences in eye and brain morphology? (ii) Does post-eclosion development shape visual performance? and (iii) Does hybridization disrupt the coordination of visual components, thereby altering performance in hybrids? Together, our findings reveal a close correspondence among post-eclosion development, behavioral maturation, and ecological divergence, illustrating how coordinated sensory evolution may contribute to reproductive isolation during speciation.

## Results

### Sympatric *Heliconius cydno* and *H. melpomene* occupy distinct visual environments

Where they coexist, *Heliconius cydno* and *H. melpomene* differ not only in warning pattern and assortative mating behavior, but also in ecology (34, 36, 47, 54). Field sampling in Colombia revealed significant differences in canopy openness between capture sites (Wilcoxon rank-sum test, W = 60, p = 0.017, r = 0.59; Fig. 1A,C), with *H. cydno* typically found in areas with higher canopy cover and *H. melpomene* in relatively more open habitats. Consistent with this, spectral irradiance differed between habitats, with closed-canopy sites exhibiting lower light intensity across wavelengths than forest-edge habitats (Fig. 1C). Because behavior also shapes the sensory input an individual receives, we also measured flight height and found that *H. cydno* flew higher on average (162.5 ± 18.7 cm SE) than *H. melpomene* (78.2 ± 26.8 cm SE; F₁,_4_₁ = 5.21, p = 0.028; Fig. 1B). These results are consistent with previous reports from Costa Rica and Panama (48–50), suggesting similar habitat differences across their shared species range. Together, these species-specific differences in habitat-use indicate that *H. cydno* and *H. melpomene* experience contrasting sensory environments, which likely impose different demands on visual processing (1–4).

**Figure 1:**
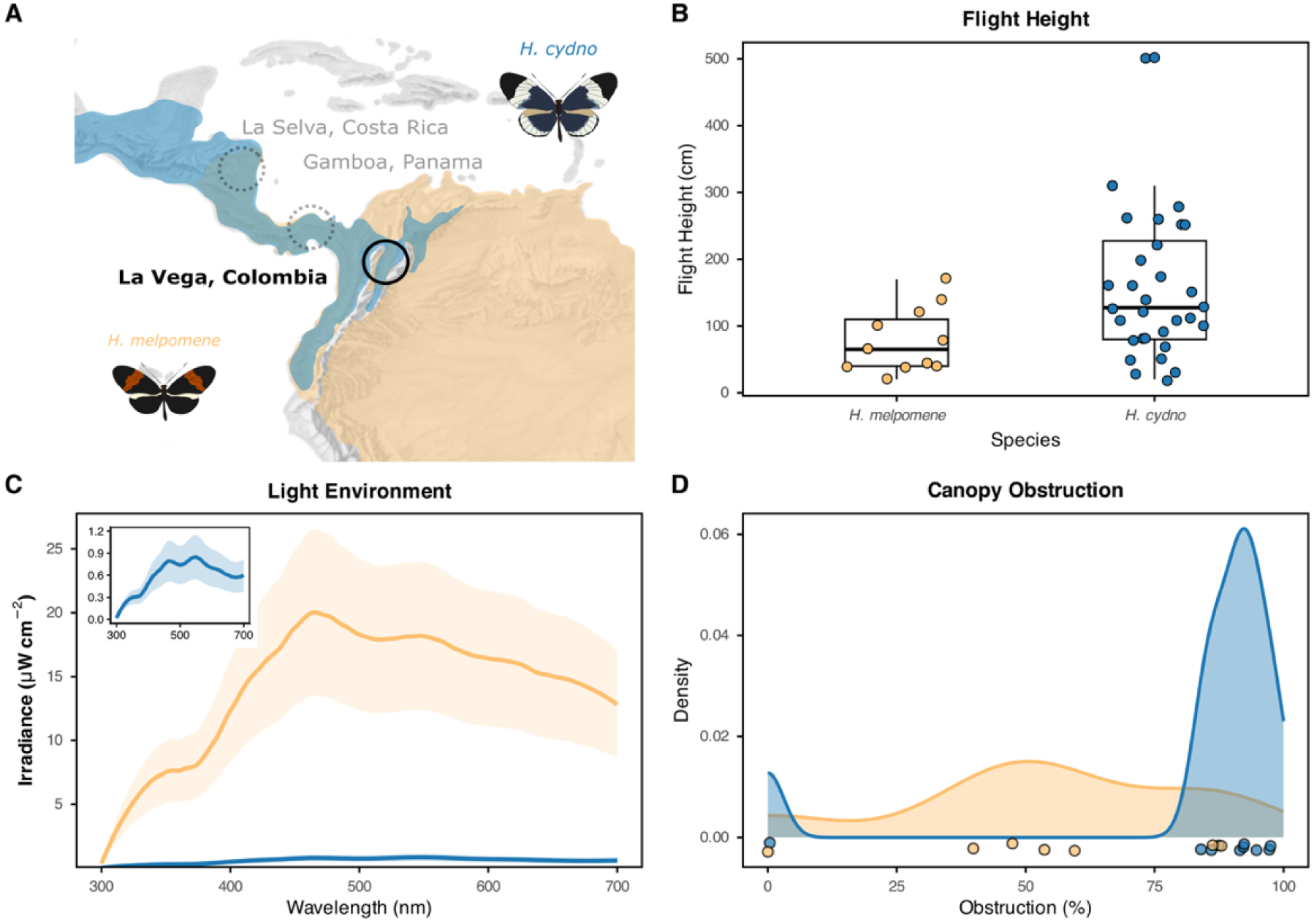
Ecological divergence between *H. cydno* and *H. melpomene*. (**A**) Geographic distribution and capture sites; black circle indicates focal population (La Vega, Colombia); dotted circles indicate previous study sites. (**B**) Flight height; boxplots show median and IQR. (**C**) Irradiance of closed-forest (blue) and forest-edge (orange) habitats; shaded areas denote SE; inset shows magnified closed-forest spectrum. (D) Density of canopy openness (%) at capture sites for *H. cydno* (blue) and *H. melpomene* (orange).

### Species, sex and maturity predict visual acuity

Given that *H. cydno* and *H. melpomene* occupy contrasting visual environments, we tested whether these ecological differences are associated with divergence in visual performance. Optomotor assays revealed clear species differences in visual acuity (Fig. 2A–B; Tables S1–S6), with *H. cydno* exhibiting higher acuity than *H. melpomene* (β = −0.080, 95% CrI = −0.108 to −0.052). Estimated mean acuity was 0.622 cpd in *H. cydno* (95% CrI = 0.548 to 0.727) and 0.480 cpd in *H. melpomene* (95% CrI = 0.426 to 0.560). Males also showed slightly higher acuity than females overall (β = 0.051, 95% CrI = 0.009 to 0.093).

**Figure 2:**
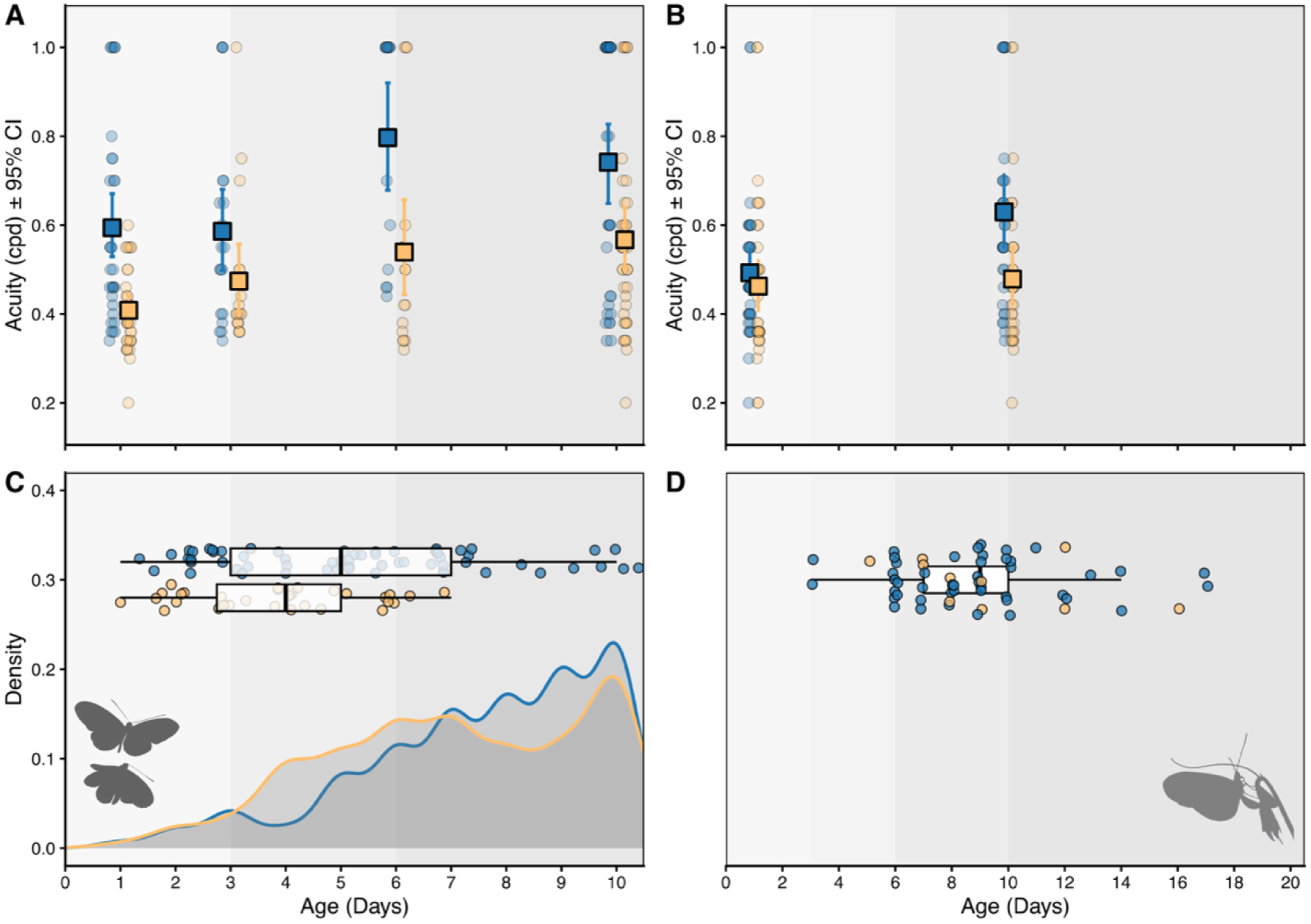
Increases in visual acuity and onset of reproductive behaviors in *H. cydno* and *H. melpomene.* **(A)** Visual acuity in males at 1, 3, 6, and 10 days post-eclosion. *H. cydno* (blue) and *H. melpomene* (orange). (**B**) Visual acuity in females at 1 and 10 days post-eclosion. Points represent individuals; large squares with error bars indicate mean 95% CI. (**C**) Onset and frequency of male courtship. Boxplots and points (top) show the distribution of ages at first courtship; density plots (bottom) illustrate the age-related increase in courtship behavior frequency across species. (**D**). Age at first oviposition for females. Boxplots and points represent individual first-laying events.

We also detected a strong and consistent effect of post-eclosion adult age on visual acuity. Across all individuals (n = 345), acuity increased with age (β = 0.012, 95% CrI = 0.008 to 0.016; Fig. 2A-B), indicating improved visual resolution during early adult life. This pattern was also evident when the species were analyzed separately. In *H. cydno* (n = 169), visual acuity increased with age (β = 0.016, 95% CrI = 0.010 to 0.022), and males again showed higher acuity than females (β = 0.085, 95% CrI = 0.016 to 0.156). Estimated mean acuity was 0.666 cpd in males (95% CrI = 0.619 to 0.723) and 0.548 cpd in females (95% CrI = 0.503 to 0.596). In *H. melpomene* (n = 176), acuity also increased with age, although more gradually (β = 0.010, 95% CrI = 0.005 to 0.014), whereas the effect of sex was weaker and its credible interval overlapped zero (β = 0.025, 95% CrI = −0.028 to 0.078). Estimated marginal means were 0.496 cpd in males (95% CrI = 0.447 to 0.569) and 0.469 cpd in females (95% CrI = 0.417 to 0.543). Together, these results show that while ecological divergence in sensory environment is associated with differences in visual acuity, early adult development also contributes to sensory maturation in both species.

### Increases in visual acuity coincide with the onset of reproductive behaviors

Male mate recognition and female oviposition in *Heliconius* are strongly visually mediated (32, 34–36), raising the possibility that improvements in visual performance coincide with the onset of reproductive maturation. To test this, we quantified male and female reproductive behaviors across the post-eclosion maturation period (Fig. S2). In standardized insectary trials, male reproductive behavior increased markedly with age, broadly overlapping with improvements in visual acuity. The onset of male courtship was strongly age-dependent, with the probability of initiating courtship increasing as males aged (Fig. 2C). Courtship began significantly earlier in *H. melpomene* (3.59 ± 1.72 days, n = 32) than in *H. cydno* (4.84 ± 2.26 days, n = 67; Weibull time ratio = 0.75, 95% CI = 0.62–0.89). Beyond onset, age also predicted courtship frequency (χ²_4_ = 293.724, p < 0.001), with older males performing courtship at higher rates. A significant age × species interaction (χ²_2_ = 8.58, p = 0.014) further indicated that this increase occurred earlier in *H. melpomene* than in *H. cydno*.

Whereas males actively search for mates, female reproductive behavior centers on host-plant selection, which requires careful visual inspection (35, 56, 57). In our insectaries, females began laying eggs around nine days post-eclosion (median = 8.5 days in *H. melpomene*, 9.0 days in *H. cydno*; Fig. 2D), with the probability of first oviposition increasing with age (Weibull shape parameter k = 2.50). Oviposition timing did not differ between species (time ratio = 0.97, 95% CI = 0.74–1.27, p = 0.82), consistent with Kaplan–Meier and log-rank tests (χ²_1_ = 0.01, p = 0.97; Fig. S3).

### *H. cydno* invests more in the eyes and optic lobe, but post-eclosion growth is restricted to the brain

Visual acuity in insects is primarily determined by the interommatidial angle, reflecting ommatidia density and number across the eye’s surface (52). Consistent with this, *H. cydno* and *H. melpomene* differed significantly in both ommatidia number and corneal area, but neither trait varied with age (Fig. 3, Tables S7–S8). *H. cydno* had more ommatidia than *H. melpomene* (χ²_1_ = 50.844, FDR-p < 0.001; Fig. 3A), and ommatidia number increased with body size (χ²_1_ = 14.198, FDR-p < 0.001), with the species difference persisting after controlling for tibia length. Age had no effect on ommatidia number (χ²_3_ = 0.997, FDR-p = 0.802; Fig. 3B). We found no evidence for an effect of sex on ommatidia number (χ²_1_ = 3.514, FDR-p = 0.162), though the smaller number of females may have limited power to detect sexual dimorphism. Corneal area showed a similar pattern, being larger in *H. cydno* and increasing with body size, but not varying with age or sex (see SI).

**Figure 3:**
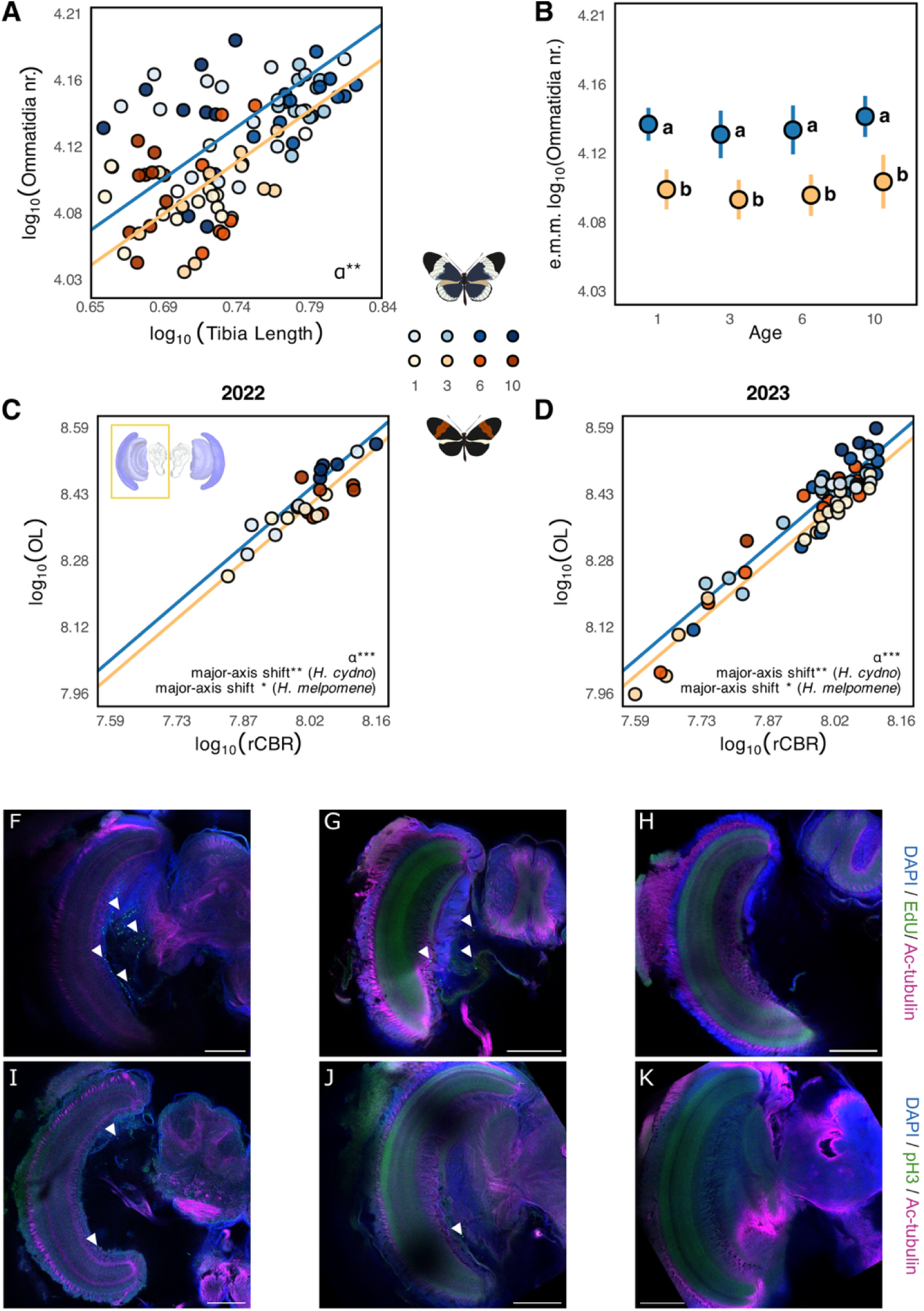
Species divergence in eye and brain morphology, and post-eclosion growth in the optic lobe. (**A**) Ommatidia number for male butterflies shows a significant elevation (α) shift, with *H. cydno* having more ommatidia for a given body size. (**B**) Estimated marginal means (e.m.m.) from linear models showing that ommatidia number remains constant across age groups (1–10 days). Letters indicate significant differences between species. (**C–D**) Standardized major axis (SMA) relationships between optic lobe (OL) and rest-of-central-brain (rCBR) volumes, showing data from both 2022 and 2023 cohorts.Interspecific divergence is characterized by elevation (α) shifts, with *H. cydno* exhibiting greater relative investment. Intraspecific maturation is dominated by major-axis shifts along common slopes, indicating coordinated post-eclosion growth. Individuals are shaded by age (light to dark) as shown in the legend. (**(F–K)** EdU **(F–H)** and pH3 (**I–K)** labeling in the optic lobe of *H. melpomene* at day 0 (F,I), 2 (G,J), and 5 (H,K) shows proliferating and mitotic cells during early adult maturation (arrowheads). All panels show merged channels: DAPI (nuclei, blue), EdU or pH3 (green), and Ac-tubulin (magenta). Scale bars: 100 µm.

To distinguish divergence along shared scaling relationships from size-independent shifts in trait investment, we used standardized major axis regression (SMATR-package; (58)). Because variance structure differed between sexes, males and females were analyzed separately (Tables S9–S10). In males, both species shared a common slope for ommatidia number against tibia length but differed in elevation (Wald χ² = 8.669, FDR-p = 0.016), indicating that *H. cydno* has more ommatidia than *H. melpomene* at equivalent body sizes (Fig. 3A). Females showed a similar pattern, although support was weaker due to smaller sample sizes (Fig. S5, Table S10).

Given pronounced species differences in eye morphology, but no age-related variation, we next asked whether this divergence extended to the brain, particularly the optic lobes. After controlling for allometric scaling relative to rest-of-central-brain volume (rCBR), SMATR analyses indicated that interspecific differences were expressed primarily as elevation shifts rather than slope changes (Fig. 3C–D; Table S12). *H. cydno* showed greater relative investment in the optic lobe overall (Wald χ²_1_ = 26.175, FDR-p < 0.001) and in four of six major optic-lobe neuropils, whereas the accessory medulla was relatively larger in *H. melpomene* (χ²_1_ = 33.155, FDR-p < 0.001). *H. cydno* also had a greater relative investment in the anterior optic tubercle (χ²_1_ = 8.122, FDR-p < 0.01) and mushroom body calyx (χ²_1_ = 7.412, FDR-p < 0.01). These results reveal pronounced species differences in sensory investment centered on the visual system, particularly the optic lobes, consistent with previous reports (6). A minor batch effect was detected, with individuals from 2022 being slightly larger than those from 2023, but overall patterns were consistent across cohorts.

Within species, age-related variation was dominated by major-axis shifts along conserved slopes, consistent with coordinated post-eclosion brain growth rather than age-specific changes in allometric scaling (Fig. 3C–D, Tables S13–S14). In *H. cydno*, significant major-axis shifts were detected in the optic lobe as a whole (Wald χ²_1_ = 16.769, FDR-p = 0.002), multiple optic-lobe neuropils, and the mushroom body (χ²_1_ = 12.591, FDR-p = 0.010). *H. melpomene* showed a similar pattern, with shifts in the optic lobe (χ²_1_ = 9.302, FDR-p = 0.038), several optic lobe neuropils, and the mushroom body (χ²_1_ = 12.950, FDR-p = 0.021). Together, these results indicate that between-species differences in brain composition reflect divergence in relative investment across visual neuropils, whereas early adult maturation within species involves coordinated expansion along conserved allometric trajectories.

### Early adult proliferation accompanies optic-lobe growth

As visual acuity and optic-lobe volumes increase during early adult life, we next tested whether these changes are supported by ongoing cell proliferation. Neural growth can arise through remodeling of existing circuits or the addition of new cells (23, 26, 59, 60). We therefore assessed proliferative activity after eclosion using EdU incorporation and anti-phospho-Histone H3 (pH3) labeling at 0, 2, and 5 days in *H. melpomene* (Fig. 3F–K). Both markers revealed a pronounced but transient period of proliferation in early adult life. At day 0, numerous EdU⁺ nuclei were detected around the optic lobes, particularly near the lobula and lobula plate (likely the inner proliferation center), with pH3⁺ cells in the same regions, indicating that cell division remains active immediately after adult eclosion. By day 2, proliferating cells were reduced and largely confined to the optic lobe, and by day 5 were no longer detected. These results indicate that proliferation declines rapidly after eclosion and coincides with the period of post-eclosion brain growth.

### F1 hybrids show intermediate visual acuity but disrupted allometric scaling and mismatched visual-system architecture

Given the pronounced post-eclosion changes in visual acuity and the possibility of disrupted development in hybrids, we assayed visual acuity in 10-day-old F1 hybrids. Acuity was intermediate between the parental species (Fig. 4A, Tables S15–S16), decreasing from *H. cydno* to F1 hybrids to *H. melpomene* (β = −0.088, 95% CrI = −0.139 to −0.036), with no deviation from a linear trend (β = −0.005, 95% CrI = −0.057 to 0.046). Estimated mean acuity declined from *H. cydno* (0.637 cpd, 95% CrI = 0.533–0.750) to F1 hybrids (0.559 cpd, 95% CrI = 0.486–0.652) to *H. melpomene* (0.479 cpd, 95% CrI = 0.404–0.562). As in the parental species, males showed slightly higher acuity than females (β = 0.067, 95% CrI = 0.010–0.124).

**Figure 4:**
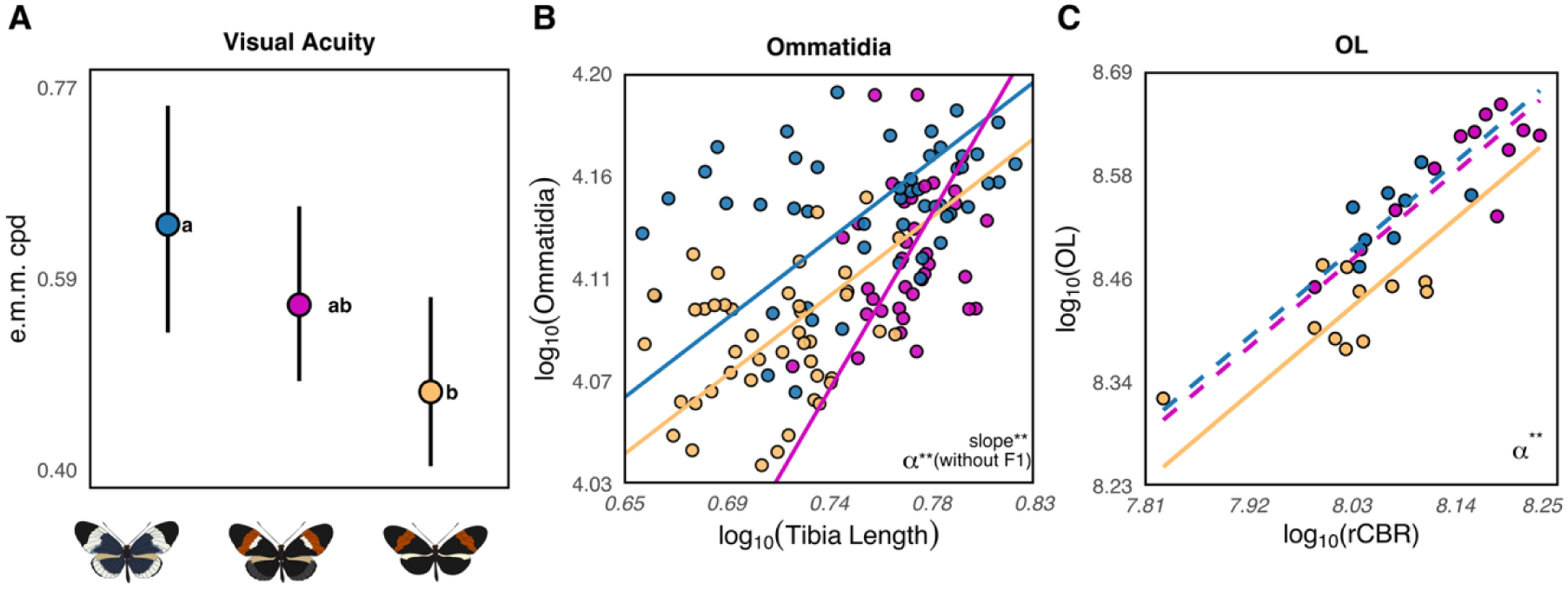
Hybrids show intermediate values of visual acuity but disrupted allometric scaling. **(A)** E.m.m. for visual acuity (cpd), in *H. cydno* (blue),F1 hybrids (magenta), and H*. melpomene* (orange). Letters indicate post-hoc group differences (**B**) Allometric relationship between ommatidia number and tibia length. Hybridization disrupts the conserved parental scaling, resulting in significant slope differences. (**C**) Scaling of optic lobe volume against rest-of-central-brain volume (rCBR) shows conserved slopes across groups, with differences expressed primarily as elevation shifts with hybrids resembling *H. cydno*.

In the subset of mature individuals for which both traits were measured (*H. cydno* = 12, *H. melpomene* = 9, hybrids = 42), facet count did not improve prediction of visual acuity beyond species and sex (Δelpd = 0.64, SE = 0.68), and its effect was highly uncertain (β = 0.88, 95% CrI = −0.66 to 2.36). This result was unchanged when including tibia length as a proxy for body size (Δelpd = −0.20, SE = 0.13). Despite intermediate visual performance, hybrid eye morphology showed complex inheritance and disrupted allometric relationships (Fig. 4B, Tables S17–S26). Ommatidia number differed among species and F1 hybrids (χ²_2_ = 55.262, FDR-p < 0.001) and increased with tibia length (χ²_1_ = 20.483, FDR-p < 0.001). Post-hoc comparisons confirmed that *H. cydno* had more ommatidia than *H. melpomene* (p < 0.001). Hybrids differed from *H. cydno* (p < 0.001), but only marginally from *H. melpomene* (p = 0.044), a difference that was not significant when restricting the analysis to males (p = 0.181; Table S18).

Major axis analyses further revealed that hybridization disrupts allometric relationships observed in the parental species (Fig. 4B). In males, slopes differed significantly between parental species and F1 hybrids for ommatidia number relative to tibia length, corneal area relative to tibia length, and corneal area relative to ommatidia number (Fig. 4B, Table S25). Pairwise comparisons showed that these differences were driven primarily by contrasts involving hybrids: *H. cydno* and hybrids differed in scaling for ommatidia number vs. tibia length (χ²_1_ = 13.429, FDR-p < 0.001), corneal area vs. tibia length (χ²_1_ = 6.594, FDR-p = 0.031), and corneal area vs. ommatidia number (χ²_1_ = 12.649, FDR-p = 0.001), while *H. melpomene* and hybrids differed in corneal area vs. ommatidia number (χ²_1_ = 12.073, FDR-p = 0.002). Although the parental species shared a common scaling relationship between ommatidia number and corneal area, this coupling was disrupted in hybrids (Fig. S10B, D). Because elevation tests assume common slopes, these differences cannot be explained as simple size-independent shifts. Instead, hybridization appears to uncouple the coordinated scaling of visual system components.

In contrast to disrupted eye allometry, optic lobe morphology in F1 hybrids retained conserved scaling and a predominantly *H. cydno*-like pattern of relative investment (Fig. 4C; Tables S27–S30). SMATR analyses controlling for rest-of-brain volume revealed no slope differences among *H. cydno*, *H. melpomene*, and hybrids for the optic lobe or individual neuropils. However, elevation shifts were detected for the optic lobe overall (Wald χ² = 12.423, FDR-p < 0.01) and most neuropils, reflecting greater relative investment in *H. cydno*, except for the accessory medulla, which was larger in *H. melpomene* (Wald χ² = 13.519, FDR-p < 0.01). Pairwise comparisons showed that hybrids generally did not differ from *H. cydno* in elevation (optic lobe: Wald χ² = 0.559, FDR-p = 1.000), but differed from *H. melpomene* (Wald χ² = 7.646, FDR-p = 0.017), supporting a largely *H. cydno*-like pattern. Where hybrids did differ from *H. cydno*, they reflected major-axis shifts along shared slopes, consistent with larger absolute brain size rather than altered proportional allocation. The lamina was an exception (Figs. S11–S12), with hybrids showing an intermediate phenotype and not differing from either species (Tables S29–S30). This likely reflects its tight coupling to the retina, as each lamina cartridge receives input from a single ommatidium (61), and ommatidia number in hybrids is shifted toward *H. melpomene*.

## Discussion

Our findings show that ecological divergence between *Heliconius cydno* and *H. melpomene* is accompanied by differences in visual performance that are further shaped by post-eclosion development. Increases in visual acuity coincide with the onset of male courtship and female oviposition –after which they are most likely to influence fitness–and post-eclosion brain growth, suggesting that early adult neurodevelopment additionally contributes to this key aspect of vision. Our results also show that first-generation hybrids exhibit intermediate visual acuity and an uncoupling of visual system components, with different traits showing contrasting patterns of inheritance. Such mismatches may alter how hybrids perceive and respond to their environment, with consequences for habitat use, behavioral performance, and gene flow.

The ecological partitioning observed at our Colombian field site mirrors that reported for other sympatric *H. cydno* and *H. melpomene* populations in Costa Rica and Panama (48–50), indicating consistent differences in visual ecology across their shared range. Closed-canopy habitats occupied by *H. cydno* are characterized by lower light intensity, altered spectral composition, and different visual backgrounds compared to the brighter, more open forest-edge environments of *H. melpomene* (51, 62). Similar ecological transitions, including differences in flight height and visual phenotypes, are seen in other *Heliconius* sister species such as *H. erato cyrbia–H. himera* and *H. erato venus–H. chestertonii*, where species in closed-forest habitats fly higher and possess more ommatidia and larger optic lobes (63–66). Together, these patterns suggest that habitat shifts repeatedly impose divergent selection on the visual system between closely related *Heliconius* taxa.

Contrasting sensory demands between closed-canopy and forest-edge habitats are expected to drive adaptive divergence in visual systems, with potential consequences for reproductive isolation when sensory traits mediate habitat use, mate recognition, or hybrid performance (4). Consistent with this, *H. cydno* and *H. melpomene* show heritable differences in visual system structure, with *H. cydno* possessing larger eyes and expanded visual brain regions, likely driven by divergent selection (6, 8). Our results provide functional support for this coordinated divergence: *H. cydno* exhibited consistently higher visual acuity than *H. melpomene* across development. This pattern aligns with broader evidence that visual performance scales with sensory investment and is shaped by the light environment and visual scene structure. For example, flight control in *Drosophila* varies with visual background (67), spiders adjust eye investment with hunting strategy and circadian niche (68) and comparative studies in fishes and birds show that acuity scales with eye size and light environment (69, 70). However, our findings further suggest that adaptive divergence is not confined to peripheral structures, but also involves coordinated changes in neural architecture that continue to mature after eclosion.

In our system, divergence in peripheral eye structure is complemented by changes in sensory processing regions of the brain (6). Compared to *H. melpomene*, *H. cydno* shows greater investment in optic-lobe neuropils involved in motion, chromatic, and polarized-light processing (71–76). Similar coordinated changes recur across ecologically divergent *Heliconius* and other insects occupying distinct light environments, including butterfly communities structured by light microhabitat and taxa diverging along diurnal and nocturnal activity niches (9, 63, 66, 77–79). Such neuroanatomical shifts are often accompanied by differences in sensory weighting, with closely related taxa differing in their reliance on visual versus olfactory cues in ways that reflect habitat-specific sensory conditions (80, 81). Consistent with this, *H. cydno* likewise relies more strongly on visual information than *H. melpomene* when positively rewarded visual and olfactory stimuli are presented in conflict (82), linking coordinated divergence across peripheral and central components to behavioral differences. At the same time, sensory systems are developmentally dynamic, and post-eclosion maturation may shape when and how these behavioral differences emerge.

Both *H. cydno* and *H. melpomene* showed coordinated volumetric growth across visual and central brain regions during early adult life, alongside improvements in visual acuity, although the timing of behavioral maturation differed. Male courtship began earlier in *H. melpomene*, indicating that similar sensory gains can follow species-specific trajectories. In *H. melpomene*, we further show that early adult brain expansion coincided with a transient period of proliferative activity around the optic lobes, with labeled cells absent by day five. Although age-related optic-lobe development has been described in other taxa (23, 24, 26–29), its behavioral consequences remain poorly understood. Our results reveal a close temporal correspondence between optic-lobe proliferation, volumetric growth, and improving visual acuity, consistent with a role for early-adult neural maturation in supporting visually mediated behavior. This window overlaps with the onset of courtship and oviposition, suggesting that sensory maturation may be critical for mate finding and host-plant use, with consequences for fitness.

Visual acuity in insects is determined by interommatidial angles, which depend on ommatidia density and thus their number for a given eye size (52). However, our developmental and hybrid data suggest a more complex picture. Rather than reflecting a single concerted developmental program, the visual system is better described as a functionally integrated mosaic, in which peripheral and central components can diverge through distinct developmental routes while remaining coupled by selection on function (14). The fact that ommatidia number remains static after eclosion, whereas acuity continues to improve, suggests that post-eclosion neural maturation and circuit refinement additionally contribute to achieved visual acuity. This integration is particularly evident in F1 hybrids, where visual traits show differing patterns of inheritance: peripheral eye traits are intermediate, or arguably *melpomene-*like, whereas optic-lobe investment is *cydno*-like, and visual acuity is intermediate. Although we cannot assign causality, these mismatches across peripheral and central components may help explain both the intermediate acuity of hybrids and the disrupted scaling relationships among visual traits.

Despite consistent habitat use and parental phenotypes, inheritance patterns observed in our Colombian populations partially differ from those reported in Panama (Fig. S10). In Panama, F1 hybrids showed *melpomene-*like ommatidia number and corneal area (8), and more intermediate, mosaic brain morphologies (6). This suggests that intraspecific variation or rearing environment may influence how visual components are integrated in hybrids. Such disrupted coordination could have evolutionary consequences. Intermediate acuity is likely suboptimal in either parental habitat, where divergent selection favors a bimodal distribution of phenotypes. Because visual performance underpins foraging, navigation, and mate recognition in *Heliconius*, these mismatched trait combinations may impose fitness costs on hybrids and contribute to reduced gene flow between these ecologically divergent species.

In summary, our results show how ecological divergence across light environments drives coordinated adaptation of the visual system, from peripheral eye morphology to central neural investment, with measurable consequences for visual performance. Although ommatidia number strongly influences visual acuity, our results suggest that it alone cannot explain this key aspect of vision. Post-eclosion brain growth, continued proliferative activity around the adult optic lobes, and age-related improvements in visual acuity, indicate that sensory maturation continues through a critical early-adult maturation window that overlaps with the onset of mate searching and oviposition. Differences in developmental timing between species further suggest that ecological adaptation shapes not only sensory phenotypes but also the developmental processes that produce them. More broadly, disruption of this coordination in hybrids generates mismatched phenotypes that are likely suboptimal in either parental environment, potentially reinforcing ecological barriers and reducing gene flow. Sensory systems thus represent not only targets of selection, but integrated developmental architectures whose integrity is essential for behavioral performance and the maintenance of species boundaries.

## Materials and Methods

Extended detailed methodology can be found in the supplementary materials.

### Animals and habitat characterization

*Heliconius cydno* and *H. melpomene* were collected near La Vega, Colombia, and reared under common-garden conditions at the Experimental Station JCM, Universidad del Rosario. F1 hybrids were generated from crosses between *H. cydno* females and *H. melpomene*. To characterize microhabitat differences between species, we sampled wild butterflies across closed-canopy and forest-edge sites near the field station. Microhabitat differences were quantified from field records of flight height at first observation following previous studies (64), and from canopy openness measured at capture locations. Additionally, we quantified the light environment by measuring downwelling irradiance (300–700 nm) in closed-forest (*H. cydno*) and forest-edge (H*. melpomene*) habitats using a spectrometer.

### Visual acuity

We quantified visual acuity in *H. cydno* and *H. melpomene* using an optomotor assay closely following Wright *et al.* 2023 (44) (see SI for details). Individuals of both sexes were tested at 1 and 10 days post-eclosion, and additional males were tested at 3 and 6 days to better resolve age-dependent changes in visual performance. F1 hybrids were also assayed at 10 days post-eclosion. Visual acuity was defined as the highest spatial frequency that elicited a consistent directional response to rotating black-and-white stripe stimuli under standardized indoor conditions.

### Onset of reproductive behaviors

To quantify age-related changes in reproductive behavior, we measured male courtship activity and the onset of female oviposition in laboratory-reared butterflies. Male courtship was assayed in standardized behavioral trials (34, 36, 55), in which individual males were paired with conspecific virgin females and scored for courtship behavior across early adult life. Female reproductive maturation was assessed as time to first oviposition by monitoring individually housed females.

### Eye- and brain-morphology and neurogenesis

Heads were dissected from freshly sampled butterflies and processed for analyses of eye and brain morphology. Eye morphology was quantified from dissected eye cuticles imaged by stereomicroscopy, from which corneal area and ommatidia number were estimated following established methods with minor modifications (83, 84). Brains were fixed in zinc-formaldehyde, stained using anti-synapsin (DSHB, RRID: AB_2315424), and imaged with confocal microscopy following previous studies (53, 85). Whole-brain image stacks were acquired, merged, and corrected for z-axis compression before volumetric analyses. Additional brains were processed for EdU incorporation and anti-pH3 staining to assess proliferative activity during early adult development, following Alcalde-Antón et al. (2023) (23, 86).

### Deep-learning-assisted brain segmentation

We measured volumes of major optic and central brain neuropils, including the lamina, medulla, lobula, lobula plate, accessory medulla, ventral lobula, antennal lobe, anterior optic tubercle, and mushroom body calyx, lobes, and pedunculus. Rest-of-central-brain volume (rCBR) was used as the allometric control, and paired structures were quantified in one hemisphere and doubled. Neuropils were segmented from confocal image stacks in Amira v.2023.2 (Thermo Fisher Scientific). To accelerate segmentation, we trained convolutional neural networks in *Biomedisa* (87, 88) to semi-automatically delineate neuropil boundaries. The training dataset included manually segmented brain volumes from 516 individuals across 43 *Heliconiini* species, compiled from previous comparative studies (18, 89, 90), and additional unpublished data. The trained networks were then applied to our dataset to generate initial segmentations, which were manually corrected in Amira to ensure anatomical accuracy. This hybrid approach substantially reduced segmentation time while maintaining consistency across individuals and brain regions.

### Statistical analyses

All analyses were performed in R v4.3.1 (91). Habitat differences were analyzed using Wilcoxon rank-sum tests and linear models. Because acuity values were bounded by the experimental setup (maximum measurable value = 1.0 cpd), visual acuity was analyzed using Bayesian censored multilevel models in *brms* (92). The timing of reproductive behaviors was analyzed using accelerated failure time models in survival (*93*) with Kaplan–Meier curves generated in *survminer* (94). Age-related variation in male mating behavior was tested using generalized linear mixed models in *glmmTMB* (95). Eye and brain morphology were analyzed using linear models (96), with Type II ANOVA in *car* (*97*) and post hoc contrasts in emmeans (*98*), together with standardized major axis regression in *smatr* (58) to assess allometric divergence among species, ages, and hybrids. Bonferroni and false discovery rate corrections were applied where appropriate. Full statistical details are provided in the Supplementary Materials.

## Acknowledgments

We thank Universidad del Rosario for insectary access at JCM and Isabel León for technical assistance. Microscopy was performed at the Center for Advanced Light Microscopy (CALM) at the LMU Munich, and at the Wolfson Bioimaging Facility, University of Cambridge, we thank these teams for their support and assistance in this work. We thank Valentin Zaitsev for helping us set up the dependencies needed to run Biomedisa.

## Funding information

Research Funded by ERC Starter Grant 851040 to R.M.M, a SWBio DTP Studentship (BB/M009122/1) to A.A.A, and a NERC IRF (NE/N014936/1) to S.H.M. Leica Stellaris 5 confocal microscope used in this study was funded by the Deutsche Forschungsgemeinschaft (DFG, German Research Foundation) – Project number 495215303.

## Notes

https://doi.org/10.5281/zenodo.19572549

### Competing Interest Statement

The authors have declared no competing interest.

