## Supplementary material for "Ecological divergence and post-eclosion brain development shape visual performance during *Heliconius* speciation": SI

**This PDF file includes:**

SI Materials & Methods
SI Results
Figures S1 to S13

Tables S1 to S30

SI References

**Other supporting materials for this manuscript include the following:**

All datasets, analysis scripts, and metadata associated with this study are publicly available on Zenodo. The full repository can be accessed at https://doi.org/10.5281/zenodo.19572549

#### Abbreviations:

CC	*H. cydno cydno*

MMC	*H. melpomene martinae*

CC x MMC	*H. cydno cydno x H. melpomene martinae* (F1 hybrids)

ME	Medulla

LAM	Lamina

LOB	Lobula

LOP	Lobula Plate

aME	accessory Medulla

vLOB	ventral Lobe of the Lobula

AOTU	Anterior Optic Tubercule

MBCA	Mushroom Body Calyx

MBPED	Mushroom Body Penduncles

MBLO	Mushroom Body Lobes

MBODY	Mushroom Body

OL	Optic Lobe

CBR	Total central brain

rCBR	total central brain minus segmented CBR neuropils

### SI Materials & Methods:

#### Study species:

We worked with *Heliconius cydno cydno* and *H. melpomene martinae* individuals collected near La Vega, Colombia, and reared under common-garden conditions at the Experimental Station José Celestino Mutis, Universidad del Rosario (5.0005° N, 74.3394° W). Outbred stocks were maintained in outdoor mesh cages (1 × 3 × 2 m) with access to 20% sugar solution, pollen sources including *Gurania*, *Lantana*, and *Psiguria* spp., and *Passiflora* spp. for oviposition. All experimental individuals were housed under common-garden greenhouse conditions following eclosion. Behavioural tests on parental individuals were conducted at 1, 3, 6, and 10 days post-eclosion to quantify age-dependent changes in visual performance. To assess whether hybrid individuals show intermediate or disrupted phenotypes, we also generated first-generation hybrids from crosses between *H. cydno* females and *H. melpomene* males. Because the strongest post-eclosion changes in visual acuity occurred during early adult life, hybrid individuals were tested at 10 days post-eclosion, when visual performance in the parental species had largely matured, allowing us to assess how visual performance and its underlying morphological components are expressed relative to the parental species.

#### Habitat characterization:

To characterize the microhabitats of *H. cydno cydno* and *H. melpomene martinae*, we conducted field sampling in the vicinity of Universidad del Rosario's José Celestino Mutis research station in La Vega, Cundinamarca, Colombia (4.955836°N, 74.381466°W). Wild individuals were captured at three ecologically distinct sites: (i) a closed-canopy temperate rainforest within the station's property (~1350 m a.s.l.; 4.955617°N, 74.379642°W), (ii) the edge of a riparian forest (~1250 m a.s.l.; 4.964609°N, 74.383777°W), and (iii) the Paraiso Andino reserve (~1500 m a.s.l.; 4.949072°N, 74.370015°W). Upon capture, each butterfly was assigned a unique identifier, and we recorded GPS coordinates, species, sex, body length (mm), and flight height before immediate release. Flight height was measured at the moment of first observation using a butterfly net handle or measuring tape, following the method described by (1). To quantify canopy cover, we took upward-facing photographs at each unique capture location, defined as non-overlapping 5 m radius zones using a Fujifilm X-Pro 3 digital camera with a 23 mm ƒ/2 lens. Photographs were taken perpendicular to the ground, with the aperture set to ƒ/8 and output in RAW format to preserve image detail. Files were initially processed in Capture One 21 and subsequently converted to binary images in ImageJ 1.53t Automated threshold analyses were applied to estimate canopy obstruction, calculated as the proportion of black pixels in the binarized image. This protocol allowed us to quantify microhabitat complexity through spatial variation in canopy openness across the different forest types occupied by the two species.

**Irradiance measurements**

Spectral irradiance was measured in representative closed-canopy (*H. cydno*) and forest-edge (*H. melpomene*) habitats using a Flame miniature spectrometer (Ocean Optics Inc.) connected to a UV–VIS optical fiber (P400-2-UV–VIS) fitted with a cosine corrector (CC-3-UV). Measurements were taken under natural light conditions between 09:00 and 12:00 to capture ecologically relevant variation in light environments. At each site, downwelling irradiance was recorded across wavelengths from 300–700 nm, with multiple replicate measurements per habitat. Irradiance values were averaged across replicates to generate habitat-specific spectra (Supplementary Fig. S13), which revealed consistent differences in both intensity and spectral composition between closed and open habitats.

#### Behavioral estimates of visual acuity:

We quantified visual acuity in *H. cydno* and *H. melpomene* using an optomotor assay following (2). Individuals of both sexes were tested at 1 and 10 days post-eclosion, with a minimum of 30 butterflies per species, sex, and age group. Because preliminary results indicated age-related improvements in visual resolution, we included additional male groups at 3 and 6 days post-eclosion to increase resolution. F1 hybrids from crosses between *H. cydno* females and *H. melpomene* males were tested at 10 days. The apparatus, following (3) consisted of a rotating wheel mounted with interchangeable vertical black-and-white stripe stimuli printed on waterproof paper (145 μm, Premium NeverTear, Xerox, CT, USA), surrounding a fixed white PVC base (33 cm diameter). The width of one cycle (a set of black and white stripes) was calculated as [(C/360)/a], where C is the circumference of the arena and a is the desired spatial frequency in cycles per degree (cpd). Butterflies were maintained in a clear Plexiglas cylinder (4 cm radius, 15 cm height) at the center of the device. All assays were conducted indoors under standardized lighting from an overhead LED ring lamp and were video recorded from above. Prior to testing, butterflies were allowed to acclimate for 5 minutes and permitted to feed on a 20% sugar water solution.

We used stimuli with spatial frequencies ranging from 0.3 to 1.0 cpd based on previous work in *Heliconius* (2), rotating alternately clockwise and counterclockwise for 10 seconds each at approximately 6 rpm. A positive optomotor response was scored only when the butterfly shifted its head or antennae in the direction of motion across two consecutive direction reversals. Stimuli were presented in ascending order of spatial frequency until no response was observed. To confirm the acuity threshold, we then tested the next two higher-frequency stimuli and again presented the last frequency that elicited a response. All responses were confirmed from video recordings, with observer identity recorded. To ensure accuracy at threshold levels, at least two independent observers confirmed the highest spatial frequency that elicited a positive response. Only individuals with clear, unambiguous behavior were retained for analysis.

#### Onset of reproductive behaviors

To quantify age-related changes in reproductive behavior, we assessed male courtship activity and the onset of female oviposition in laboratory-reared butterflies.

Male mating behavior was measured using 15-minute trials in which individual males were placed in a mesh cage (1 × 3 × 2 m) with a conspecific virgin female (0–4 days post-eclosion), following established protocols (4–6). Males were tested between 1 and 10 days post-eclosion (10:00–16:00) under natural light and allowed to acclimate for one hour prior to trials. Trials were divided into one-minute intervals, and courtship behavior (sustained hovering or chasing) was scored as present or absent per interval; repeated courtship within a minute was recorded once, but behaviors spanning multiple minutes were scored in each interval. Weather conditions were recorded at the start of each trial (overcast, cloudy, or sunny), and trials were not conducted during rain. If copulation occurred, individuals were gently separated to prevent mating, a procedure shown not to affect subsequent behavior (5, 7). Females that did not mate during a trial were reused in other tests conducted on the same day.

Female reproductive maturation was assessed by measuring time to first oviposition. Females (47 *H. cydno*, 10 *H. melpomene*) were housed individually in mesh cages provisioned with *Passiflora* spp. and monitored daily from eclosion. We recorded species, date of eclosion, mating date (inferred from spermatophore presence or direct observation), and the date of first confirmed egg laying. Only verified oviposition events were used to score reproductive onset.

#### Eye and brain sampling and preparation

Complete heads were removed from freshly sampled butterflies, fixed in zinc-formaldehyde solution (ZnFA; 0.25% ZnCl₂, 0.788% NaCl, 1.2% sucrose, 1% formaldehyde) for 16–20 hours at room temperature under gentle agitation, and then incubated in 80% methanol / 20% DMSO for 2 hours. Samples were subsequently transferred to 100% methanol and stored at −20 °C until processing.

Eye morphology analysis followed previously published methods (8, 9), with minor modifications. In brief, specimens were thawed at room temperature, and both eyes were removed and incubated in 20% sodium hydroxide (NaOH) for 18–24 hours. Each eye cuticle was subsequently cleaned of excess tissue, mounted on a microscope slide in Euparal (Carl Roth GmbH), and left to dry overnight before imaging. Slides were then imaged at 25× magnification using a Leica M80 stereomicroscope fitted with a Leica Flexacam C1 camera and Leica Application Suite X software. Eye surface area was measured using the Freehand Selection and Measure tools in ImageJ/Fiji (10), while total ommatidia (facet) number was quantified using image thresholding and the Analyze Particles function. To account for body size variation, inter-ocular width and hind tibia length were measured using the Straight Line and Measure tools. The left eye was used for subsequent analyses unless damaged, missing, or of poor image quality, in which case the right eye was substituted.

Brain morphology was visualized using immunofluorescence staining following established protocols (11, 12), with minor modifications. Brains were rehydrated through a graded methanol series (90%, 70%, 50%, 30%, and 0%, each in 0.1 M Tris buffer, pH 7.4; 10 minutes per step). Tissues were pre-incubated for 2 hours at room temperature in 5% normal goat serum (NGS) diluted in PBS containing 1% DMSO and 0.005% sodium azide (PBSd-NGS). Primary antibody 3C11 (anti-synapsin, Developmental Studies Hybridoma Bank, RRID: AB_2315424) was applied at a 1:30 dilution in PBSd-NGS and incubated for 3.5 days at 4 °C under continuous agitation. Following three 2-hour rinses in PBSd, samples were incubated with a Cy3-conjugated goat anti-mouse IgG secondary antibody (Jackson ImmunoResearch; Cat. No. 115-165-146; RRID: AB_2338690), diluted 1:100 in PBSd-NGS, for 2.5 days at 4 °C under agitation. After staining, samples were dehydrated through an ascending glycerol series (1%, 2%, 4% for 2 hours each, followed by 8%, 15%, 30%, 50%, 60%, 70%, and 80% for 1 hour each) in 0.1 M Tris buffer with 1% DMSO, then transferred to 100% ethanol and cleared in methyl salicylate.

Confocal imaging was performed using a Stellaris 5 confocal laser-scanning microscope (Leica Microsystems, Mannheim, Germany) with a 10× dry objective (NA 0.4; Leica Material No. 11506511), a mechanical z-step of 2 μm, and an x–y resolution of 512 × 512 pixels. Whole-brain stacks were acquired as image stacks (20% overlap) which were automatically merged. The z-dimension was scaled by a factor of 1.52 to correct for artifactual compression associated with the use of a 10× air objective (13).

#### Neurogenesis: EdU and anti-pH3 staining

To assess patterns of neurogenesis, we stained brains from *H. melpomene* collected at 0, 2, and 5 days post-eclosion. These time points were selected to precede visual acuity assays conducted at 1, 3, 6, and 10 days, based on the hypothesis that proliferative activity may precede functional changes in visual processing. We applied two complementary approaches, immunofluorescence staining for phosphorylated histone H3 (pH3), a marker of cells undergoing mitosis (14, 15) and EdU incorporation, which labels cells in S-phase, following established protocols (16, 17). Brains were prepared and fixed as described above for synapsin immunohistochemistry. For pH3 staining, whole heads were fixed in ZnFA and stored in methanol. After rehydration, samples were incubated with a rabbit anti-phospho-Histone H3 (Ser10) (Alexa Fluor® 647 conjugate) antibody (Cell Signalling, #3458) at 1:500 dilution and a mouse anti-acetylated tubulin (Merk, #T7451) at 1:150 in PBSd-NGS for 3.5 days at 4 °C under gentle agitation (17). The rest of the protocol was the same as the one described above, with the addition of an extra step. After the secondary antibody incubation brains were washed in PBSd (3 × 30 min) and in H_2_O with 0.2% Triton (1× 30 min), and then they were counterstained with DAPI (Sigma-Aldrich, #D9542; 1:1000 in H_2_O with 0.2% Triton) for 3 h at room temperature to visualize cell nuclei.

#### For EdU labelling, we followed a modified protocol for large insect brains (17). Briefly, a small window was cut in the head capsule of anesthetized adults, and intact heads were incubated for 3 h in 20 μM EdU diluted in Grace’s Medium (Thermo Fisher, #11595030). Brains were then dissected and fixed in in ZnFA overnight at 4 °C. After permeabilization in PBS with 1% Triton, incorporated EdU was detected using the Click-iT EdU Imaging Kit (Thermo Fisher, #C10420) following manufacturer’s instructions. Following the Click-iT reaction, immunohistochemistry was performed using anti-acetylated tubulin as the primary antibody and DAPI as a nuclear marker. Following both protocols, tissues were clarified using a graded glycerol and ethanol dehydration series, cleared in methyl salicylate, and imaged using the same confocal microscopy parameters described above.

#### Brain segmentation

We measured volumes of key optic and central brain regions, including the lamina, medulla, lobula, lobula plate, accessory medulla, ventral lobula, antennal lobe, anterior optic tubercle, and mushroom body calyx, lobes and pedunculus. The rest-of-central-brain (rCBR) was used as an allometric control. Paired structures were quantified in one hemisphere and doubled. Brain regions were segmented from confocal image stacks using Amira v.2023.2 (Thermo Fisher Scientific). To accelerate segmentation, we trained convolutional neural networks using *Biomedisa* (18, 19) to semi-automatically delineate neuropil boundaries. The training dataset included manually segmented brain volumes from 516 individuals across 43 Heliconiini taxa. Of these, 140 individuals representing 31 species were derived from previously published datasets (20–22) while an additional 376 individuals representing 5 species and 7 hybrid groups were obtained from unpublished data (including this study). Together, this combined dataset provides a broad representation of interspecific and hybrid variation in brain morphology for training the convolutional neural networks. The trained networks were applied to our dataset to generate initial segmentations, which were manually refined in *Amira* to ensure anatomical accuracy. This hybrid approach substantially reduced annotation time while maintaining consistency across individuals and brain regions.

#### Biomedisa training workflow

#### We trained convolutional neural networks using the offline version of Biomedisa (18, 19), which employs a three-dimensional U-Net architecture optimized through Keras and TensorFlow. All training and validation were performed on the high-performance computing cluster of the LMU Faculty of Biology, equipped with four NVIDIA A100-SXM4-80 GB GPUs.

Training images and corresponding label stacks were organized into separate directories for training and validation. Networks were trained directly on full three-dimensional image volumes, with a stride size of 16 voxels to generate overlapping sampling regions across the dataset. Models were trained for up to 80 epochs with a learning rate of 0.001 and a batch size of 24, and validation performance was assessed every two epochs.

To improve generalization, data augmentation was applied by flipping along the y-axis and randomly rotating each stack by up to 25°. The parameters --acwe_smooth 100, --clean 0.1, and --fill 0.9 controlled contour smoothing and the removal or interpolation of small artifacts during segmentation refinement. The --only flag specified the subset of neuropil labels included in the training dataset. An early stopping criterion was implemented if validation accuracy failed to improve within eight consecutive checks.

Model accuracy was evaluated using the Dice similarity coefficient, which measures overlap between predicted and manually annotated segmentations. After training, the resulting multi-label 8-bit TIFF stacks were exported and visually inspected in *Amira* v.2023.2. Minor boundary inaccuracies were corrected before volumetric quantification to ensure consistency across all samples.

#### Statistical analysis:

##### Habitat characterization and flight height

To test whether canopy obstruction differed between species, we used a non-parametric Wilcoxon rank-sum test on canopy cover estimates from capture locations. Effect sizes were quantified as rank-biserial correlations. To test for species differences in flight height, we fitted a linear model (lm function, base R) with species as the predictor. Site and sex were initially included but did not improve model fit, and the random effect of site (tested with lmer in the lme4 package) collapsed to zero variance, so they were excluded.

##### Behavioral estimates of visual acuity

### We analyzed how species identity, sex, and post-eclosion age influence visual acuity in *H. cydno* and *H. melpomene*. Because acuity values were bounded by the experimental setup (maximum measurable value = 1.0 cpd), we implemented Bayesian censored multilevel models fitted (brms package (23). Visual acuity (log10-transformed) was modeled as a function of species, age, and age, and sex, with observer identity and individual ID were included as random intercepts to account for repeated scoring and individual-level variation. Censoring was specified for individuals with acuity values at the upper measurement limit (1.0 cpd). We also tested alternative formulations including a species-by-age interaction, a species-by-sex interaction, and testing temperature as additional predictors, but these did not improve predictive performance based on approximate leave-one-out cross-validation (LOO), so the simpler model was retained. Weakly informative priors were chosen to reflect plausible log10-transformed response values: normal (0, 1) for fixed-effect slopes, normal (–0.2, 1) for the intercept, and exponential (1) for residual and random-effect standard deviations. Models were fitted using four chains (6,000 iterations; 3,000 warm-up) with conservative sampling parameters (adapt_delta = 0.9999, max_treedepth = 15) to ensure stable sampling and minimize divergent transitions. Convergence and sampling performance were evaluated using R-hat, effective sample sizes, pairwise parameter checks, and posterior predictive checks.

##### Onset of reproductive behaviors

To quantify the timing of reproductive maturation, we analysed the age at onset of female oviposition and male courtship using accelerated failure time (AFT) models. For male courtship onset, age at first observed courtship was treated as a time-to-event variable. For females, time to first confirmed oviposition was analysed similarly. In both cases, all individuals experienced the event during the observation period, and no censoring was included.

We fitted exponential and Weibull AFT models with species as a factor using the *survival (24)* . package in *R*. The exponential model assumes a constant hazard with age, whereas the Weibull model allows the hazard to increase or decrease over time. Models were compared using likelihood-ratio tests, and the Weibull distribution was selected where it provided a significantly better fit. From Weibull models, we estimated the shape parameter (k), which characterizes whether the probability of behavioral onset (courtship or oviposition) accelerates with age, and calculated species-specific time ratios with 95% confidence intervals.

For visualization, Kaplan–Meier curves were generated using *survminer* (25), showing the proportion of individuals that had not yet initiated courtship or oviposition as a function of age, together with 95% confidence intervals and risk tables.

##### Effect of age on mating behavior

For male mating behavior, we obtained count data representing the number of one-minute intervals in which a male performed a given behavior during each trial. Because males were observed repeatedly across post-eclosion ages, and trials were nested within cages and involved different females, we used mixed-effects models to account for this hierarchical structure. We analyzed the effect of male age on mating behavior using generalized linear mixed models (GLMMs) fitted with the package glmmTMB (26) in R.

Separate models were fitted for courtship behavior, mating attempt, and total mating behaviors. Male age was included as the main predictor and modelled either as a linear term or as a quadratic polynomial (raw scale) when visual inspection suggested non-linearity. Species identity and weather conditions were included as fixed effects. Random effects for male identity, female identity, and cage were initially specified, and the random-effects structure was simplified where necessary based on Akaike Information Criterion (AIC) (27). Behavioral counts were modelled using a negative binomial distribution with the nbinom1 parameterization, which assumes variance increases linearly with the mean. Fixed effects were assessed using likelihood-ratio tests, and model diagnostics were evaluated using DHARMa (28). Species- and age-specific predictions were extracted using emmeans (29) for visualization.

##### Eye morphology

Analyses of eye morphology focused on corneal surface area and total ommatidia (facet) number. Both traits were log_10_-transformed prior to analysis. To account for body size variation, log_10_-transformed hind tibia length and log_10_ interocular width were included as covariates. Linear models were fitted with species, sex, and age as fixed effects, and interaction terms were sequentially removed based on likelihood ratio tests. Fixed effects were evaluated with Type II ANOVA (*car (30)*), and model residuals were inspected for normality. Estimated marginal means (*emmeans (29)*) were extracted for significant predictors. To complement linear modelling, standardized major axis (SMA *(31)*) regression was used to test allometric relationships between log_10_ area and log_10_ tibia length, and between log_10_ tibia length and log_10_ ommatidia count. For each comparison, common slopes were estimated, and species- or age-specific differences were evaluated as slope differences, elevation shifts (intercept differences at a common slope), or major axis shifts (parallel displacements along a common slope). Model fit was evaluated with likelihood ratio tests, and effect sizes were expressed as correlation coefficients derived from test statistics. Results from both linear models and SMA analyses are summarized in Tables S1-S4.

##### Brain morphology

All analyses were performed in R v4.3.1 (32). For each neuropil, we fitted Gaussian linear models with log_10_-transformed neuropil volume as the response variable and included log_10_-rest-of-central-brain volume (rCBR), species, age, and their interaction (species × age) as fixed effects. Interaction terms were retained when supported by likelihood-ratio tests on nested models (drop1, test = "Chisq") and removed otherwise to obtain minimal adequate models. Fixed effects were evaluated using Type II ANOVA implemented in the car package. Model assumptions were assessed by inspection of Q–Q plots, residuals-versus-fitted plots, and residual histograms, and influential observations were examined using Cook’s distance. For significant predictors, estimated marginal means and pairwise contrasts were obtained using the emmeans package (29), with with Bonferroni correction applied to account for multiple comparisons. Results from the linear models are summarized in Table S11 for comparisons between *H. cydno* and *H. melpomene* across ages (n = 92), and in Table S21 for comparisons between parental species and F1 hybrids (n = 34), with Bonferroni-corrected pairwise comparisons reported in Table S22.

Standardized major axis (SMA) regressions were conducted in parallel using the smatr package (31), following established approaches in *Heliconius* neuroanatomical studies (12). For each neuropil, we first confirmed a significant allometric relationship between log_10_-neuropil volume and log_10_-rCBR. We then tested for group differences using a sequential procedure: (i) heterogeneity in allometric slopes among groups, (ii) if slopes were homogeneous, differences in elevation (intercept) along a common slope, and (iii) major-axis (grade) shifts along that common slope. Species was used as the grouping factor for analyses of the combined dataset, whereas age was used as the grouping factor in species-specific subsets to assess whether early adult brain growth produced coordinated scaling across neuropils within each species. Additional SMA analyses including F1 hybrids were conducted to evaluate their position relative to the parental species, and pairwise SMA comparisons were performed where appropriate. Robust estimation was enabled for all SMA analyses (robust = TRUE). We report test statistics, p values, and effect sizes as correlation coefficients (r) derived from the reported statistics; for elevation and major-axis shift tests, direction indices (DI) are also provided where applicable. When summarizing SMA outcomes across multiple neuropils, p values for slope, elevation, and major-axis shift tests were adjusted using the Benjamini–Hochberg false discovery rate (FDR) procedure. SMA results for interspecific comparisons and within-species age analyses are presented in Tables S12–S14. SMA analyses including hybrids are reported in Table S23, with corresponding pairwise comparisons in Table S24. Supplementary figures illustrating scaling relationships are provided in Figs. S6–S8 and S11–S12.

SI Results:

#### Effect of age on mating behaviors

The frequency of male chasing behavior was strongly influenced by age, species, and weather, with a significant age × species interaction (LRT: χ²₂ = 8.90, p = 0.012). Removing age from the model resulted in a substantial loss of fit (χ²₄ = 271.54, p < 0.001, ΔAIC = 263.5), confirming that age is the primary predictor of variation in chasing. Species identity also contributed significantly (χ²₃ = 14.50, p = 0.0023), indicating overall differences in chasing rates between *H. cydno* and *H. melpomene*. Weather had a strong effect as well (χ²₂ = 30.93, p < 0.001), with males chasing more frequently under sunnier conditions. Together, these results show that chasing increases with male age in both species, but the trajectory differs significantly between them, and environmental light conditions further modulate this courtship component.

Hovering behavior was strongly influenced by male age, with model fit declining substantially when age was excluded (LRT: χ²₂ = 264.9, p < 0.001, ΔAIC = 260.9). Species identity also contributed significantly (χ²₁ = 12.1, p < 0.001, ΔAIC = 10.1), indicating consistent differences between *H. cydno* and *H. melpomene*. Weather conditions had an additional effect (χ²₂ = 27.7, p < 0.001, ΔAIC = 23.7), with hovering occurring more frequently under sunnier conditions. Together, these results demonstrate that hovering, like chasing, increases with male age but is further modulated by both species identity and environmental light conditions.

The frequency of mating attempts was strongly predicted by male age, with model fit declining sharply when age was excluded (LRT: χ²₂ = 196.5, p < 0.001, ΔAIC = 192.5). Species identity also had a weaker but significant effect (χ²₁ = 3.95, p = 0.047, ΔAIC = 2.0), indicating differences between H. cydno and H. melpomene. Weather conditions further explained variation (χ²₂ = 24.3, p < 0.001, ΔAIC = 20.3), with mate attempts occurring more frequently under sunnier conditions. These results show that mate attempts, like chasing and hovering, increase markedly with age and are further modulated by species identity and environmental conditions.

Guarding behavior was primarily driven by male age, with model fit deteriorating substantially when age was excluded (LRT: χ²₂ = 129.8, p < 0.001, ΔAIC = 125.8). Species identity also explained significant variation (χ²₁ = 4.43, p = 0.035, ΔAIC = 2.4), suggesting that *H. cydn*o and *H. melpomene* differ in the propensity to guard. These results indicate that guarding, like other courtship components, increases with male age and also varies between species.

Total mating behaviors were overwhelmingly predicted by male age, with model fit collapsing when age was excluded (LRT: χ²₂ = 290.5, p < 0.001, ΔAIC = 286.5). Species identity also explained significant variation (χ²₁ = 8.49, p = 0.0036, ΔAIC = 6.5), reflecting differences between H. cydno and H. melpomene. Weather conditions further contributed to variation (χ²₂ = 27.0, p < 0.001, ΔAIC = 23.0), with higher overall activity under sunnier conditions. These results confirm that the cumulative expression of mating behaviors increases sharply with age and is further modulated by both species identity and environmental conditions.

Figures:

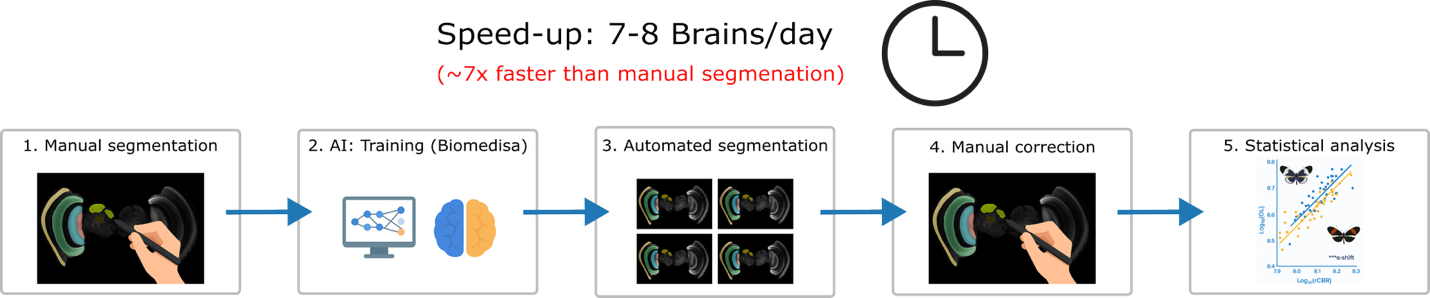

**Fig. S 1: Workflow for large-scale brain segmentation and analysis using Biomedisa pipelines.**
A streamlined five-step pipeline used to quantify Heliconius brain neuropils at scale. (1) A subset of brains is manually segmented to generate high-quality training data. (2) These segmentations are used to train a Biomedisa neural network. (3) The trained model performs automated segmentation on full datasets, achieving a 7–8× speed-up relative to fully manual workflows. (4) Automated outputs are manually checked and corrected to ensure anatomical accuracy. (5) Final volumetric datasets are used for statistical analyses of brain morphology and allometric scaling.

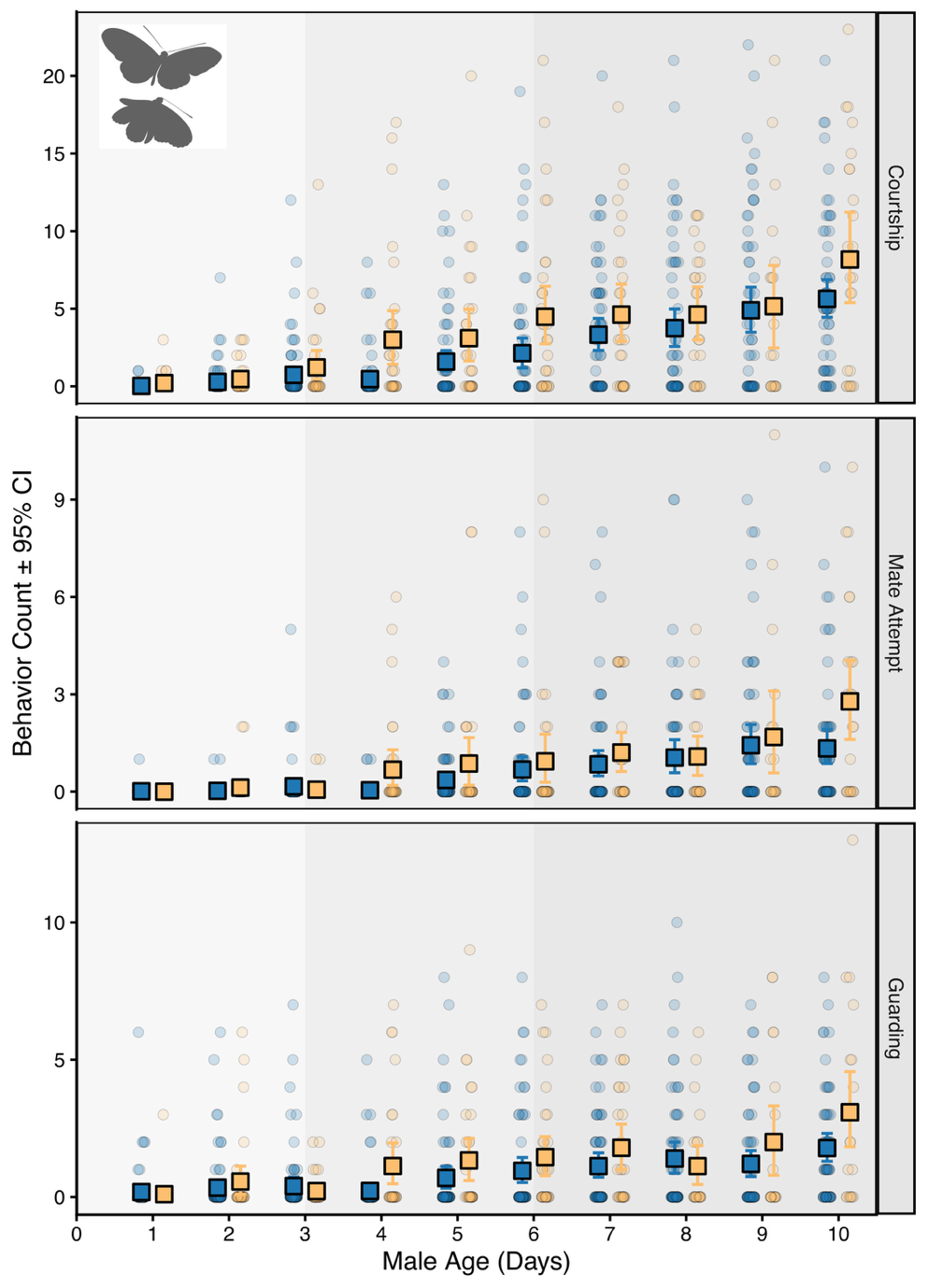

**Fig. S 2:** **Age-related changes in mating behaviours, in H. cydno (blue) and H. melpomene (orange).** Age-related increase in male courtship behaviours.

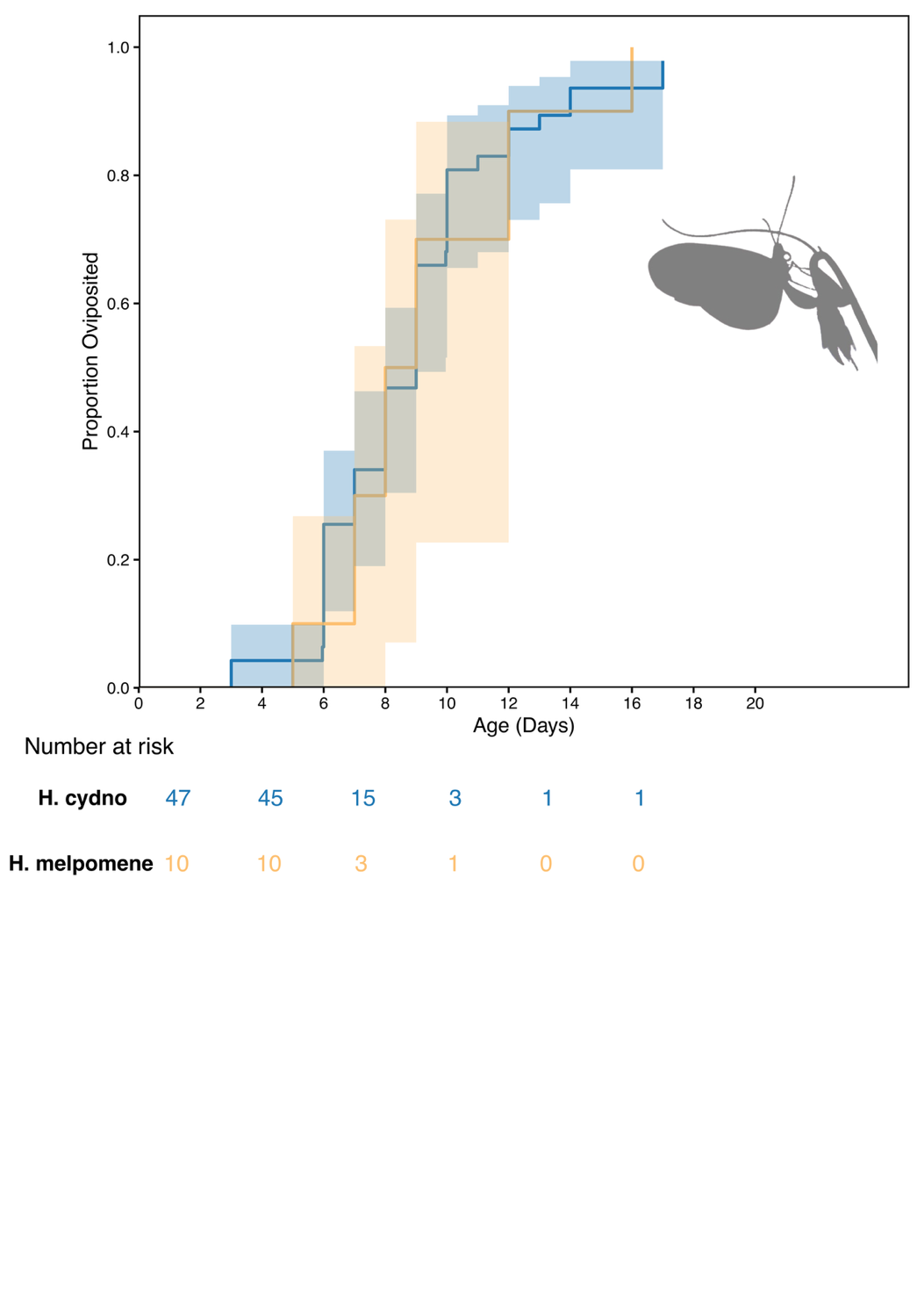

**Fig. S 3:** **Age at first oviposition in *H. cydno* and *H. melpomene*.** Kaplan–Meier survival curves showing the proportion of females that had not yet oviposited across age. Shaded areas represent 95% confidence intervals. H. cydno (blue) and H. melpomene (orange) show broadly similar timing of reproductive onset, with most individuals initiating oviposition between 5–15 days post-eclosion. Numbers at risk are shown below.

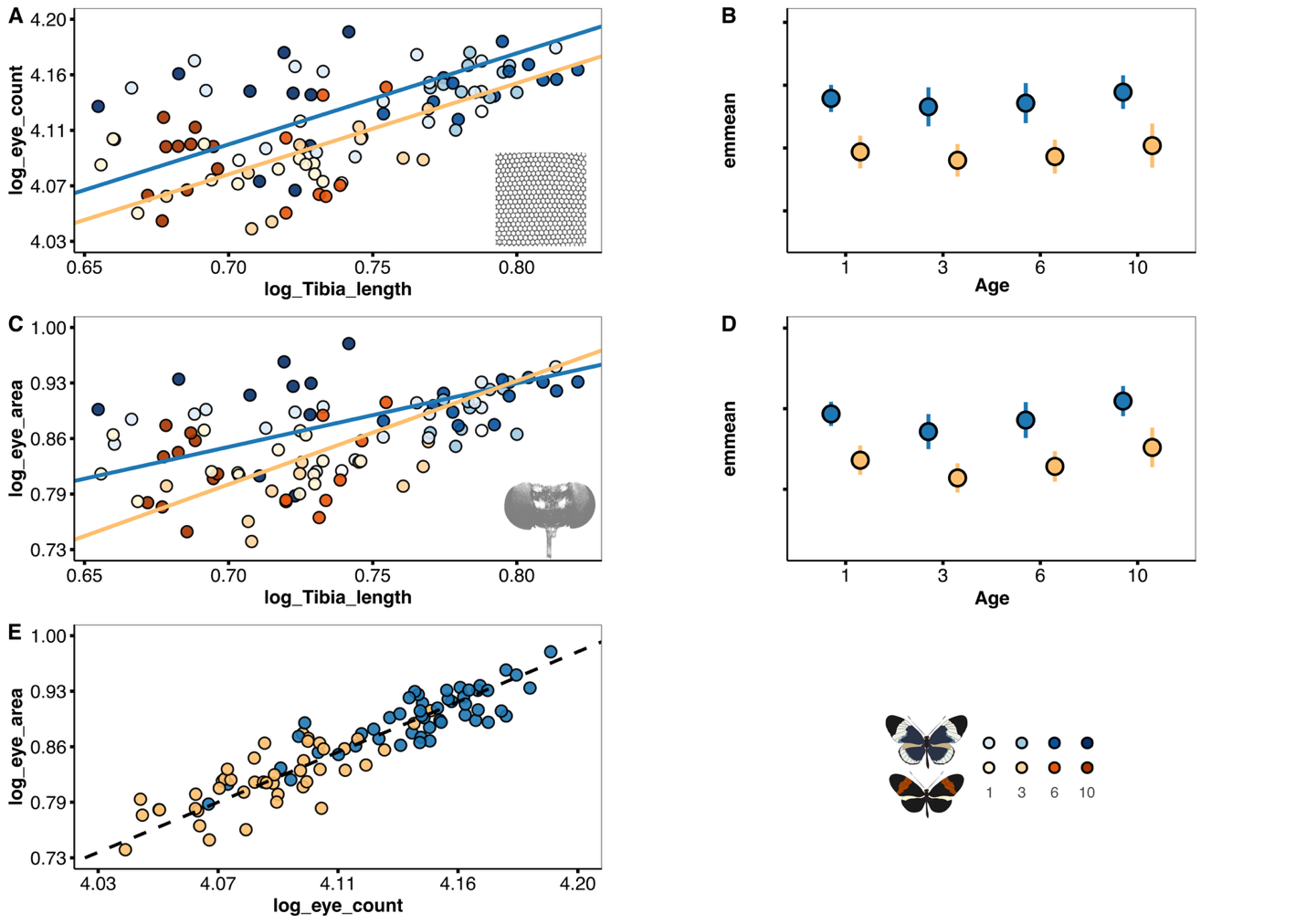

**Fig. S 4: Species differences in eye morphology between male H. cydno (blue) and H. melpomene (orange**). (**A**) Facet count against tibia length shows a significant elevation (α) shift, with H. cydno having more facets for a given body size. (**B**) Estimated marginal means (EMMs) for facet count across age (1, 3, 6, and 10 days) demonstrate that facet count remains constant with age but differs significantly between species. (**C**) Total eye area against tibia length similarly shows an elevation shift, with H. cydno maintaining larger eye areas across the sampled body size range. (**D**) EMMs for eye area across age groups. (E) Eye area against facet count shows a strong correlation along a common major-axis slope (dashed line), suggesting that the increased facet count in H. cydno is primarily driven by an overall increase in eye size. In scatterplots (A, C), individual age is represented by a color gradient (light to dark), and solid lines represent predicted allometric relationships. Error bars in (B, D) represent 95% credible intervals.

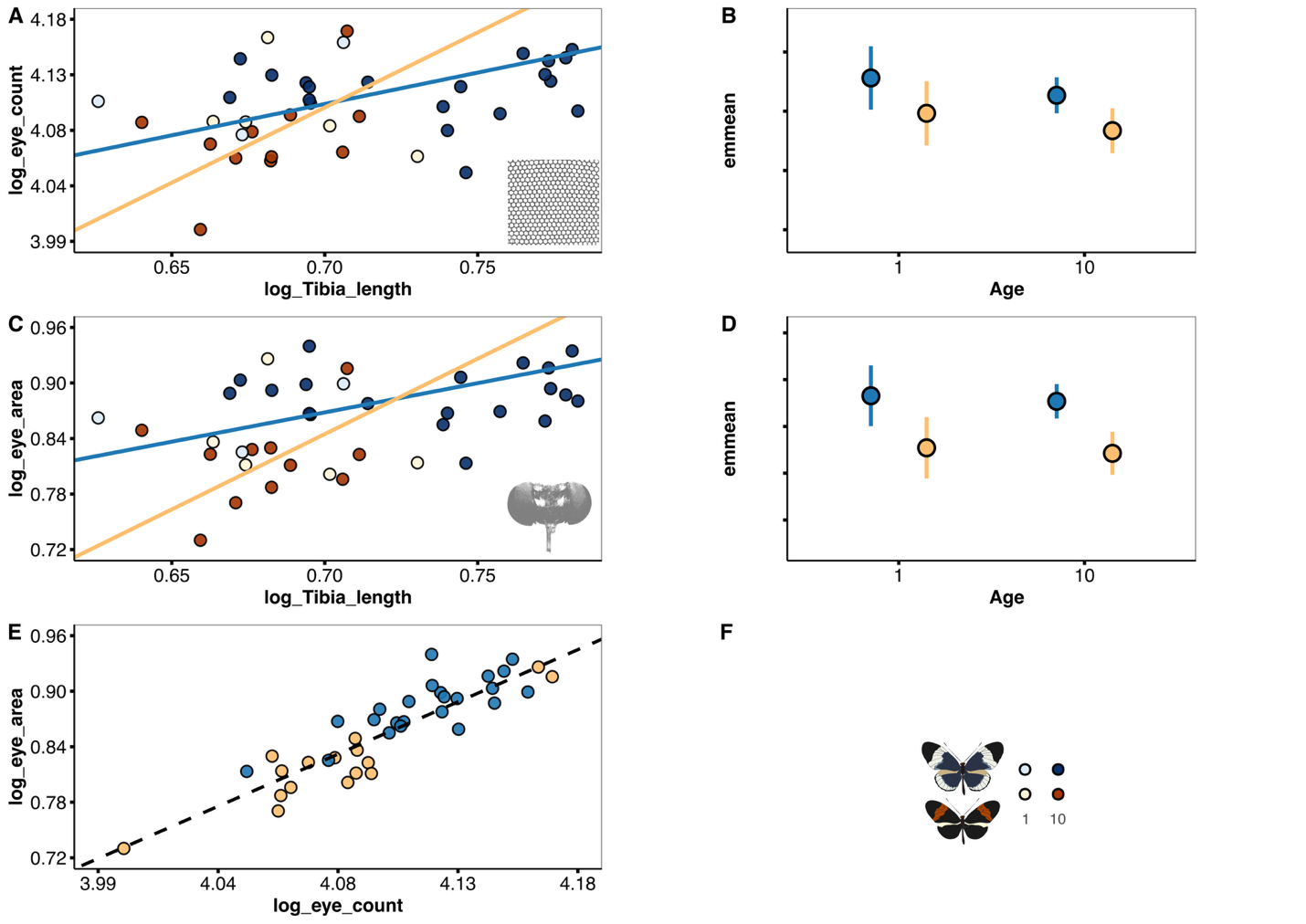

**Fig. S 5: Species differences in eye morphology between female H. cydno (blue) and H. melpomene (orange**). (**A**) Facet count against tibia length and **(C**) total eye area against tibia length. While H. cydno generally exhibits larger values for a given body size, the apparent differences in allometric slopes between species in these panels are likely driven by reduced statistical power due to the smaller female sample size relative to males. (**B, D**) E.M.M.s for facet count and eye area across age (1 and 10 days), showing stable species differences across the sampled ages. (**E**) Eye area against facet count shows a strong correlation along a common major-axis slope (dashed line). In scatterplots (A, C), individual age is represented by color saturation (light = 1 day, dark = 10 days), and solid lines represent predicted allometric relationships. Error bars in (B, D) represent 95% credible intervals.

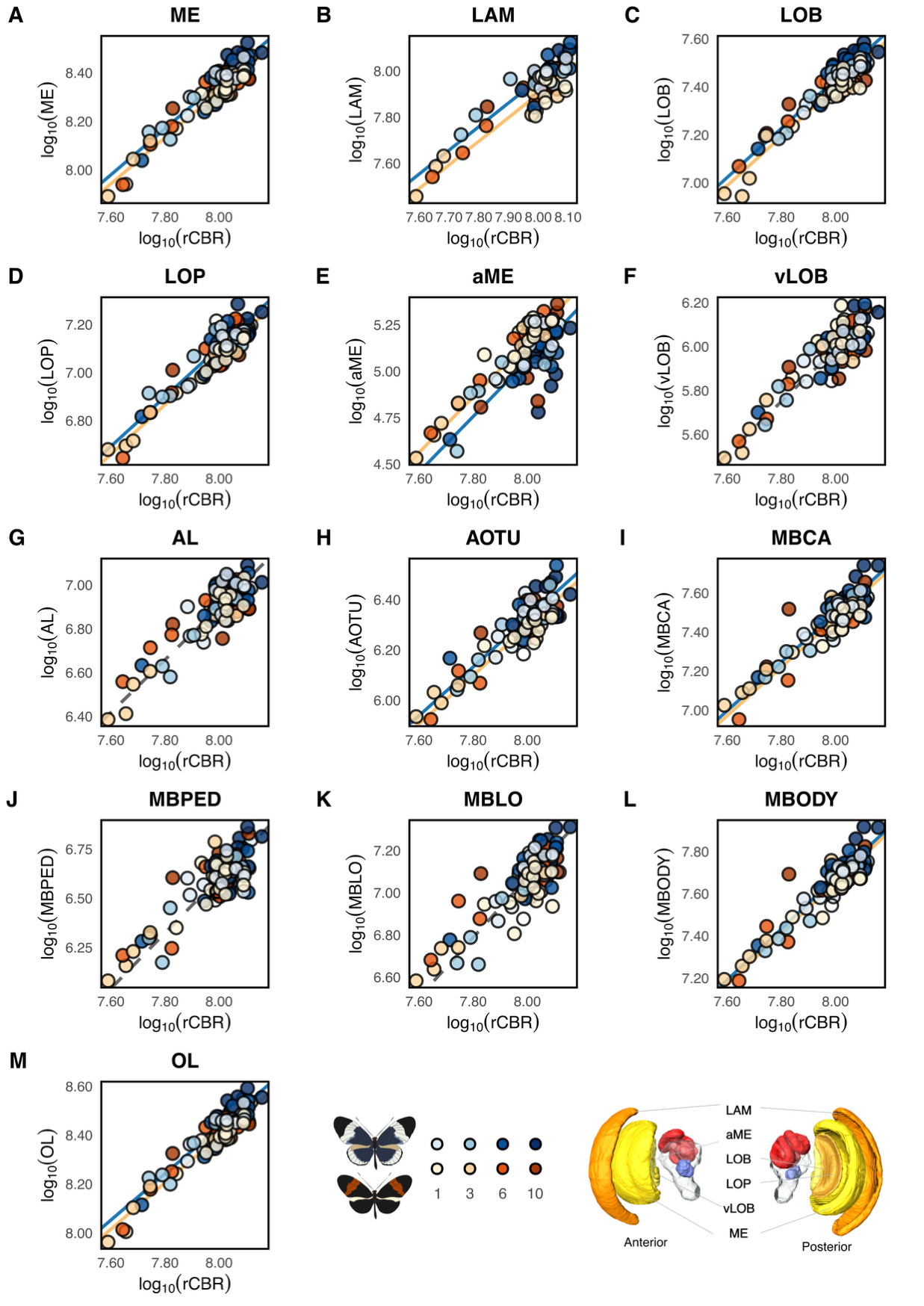

**Fig. S 6**: **Divergence and allometric relationships across visual neuropils in** **H. cydno** **and** **H. melpomene**. **(A–M)** All traits are log10-transformed. Scatterplots show the relationships between individual neuropil volumes and the rest-of-central-brain volume (rCBR). Lines indicate species-specific Standardized Major Axis (SMA) regressions (solid) or common slopes where no significant difference in slope was detected (dashed). Interspecific divergence is primarily characterized by elevation (α) shifts, with H. cydno generally exhibiting greater relative investment in visual neuropils. Intraspecific maturation is characterized by shifts along common major-axis slopes, indicating coordinated post-eclosion growth. Individuals are shaded by age from 1 to 10 days (light to dark) as shown in the legend.

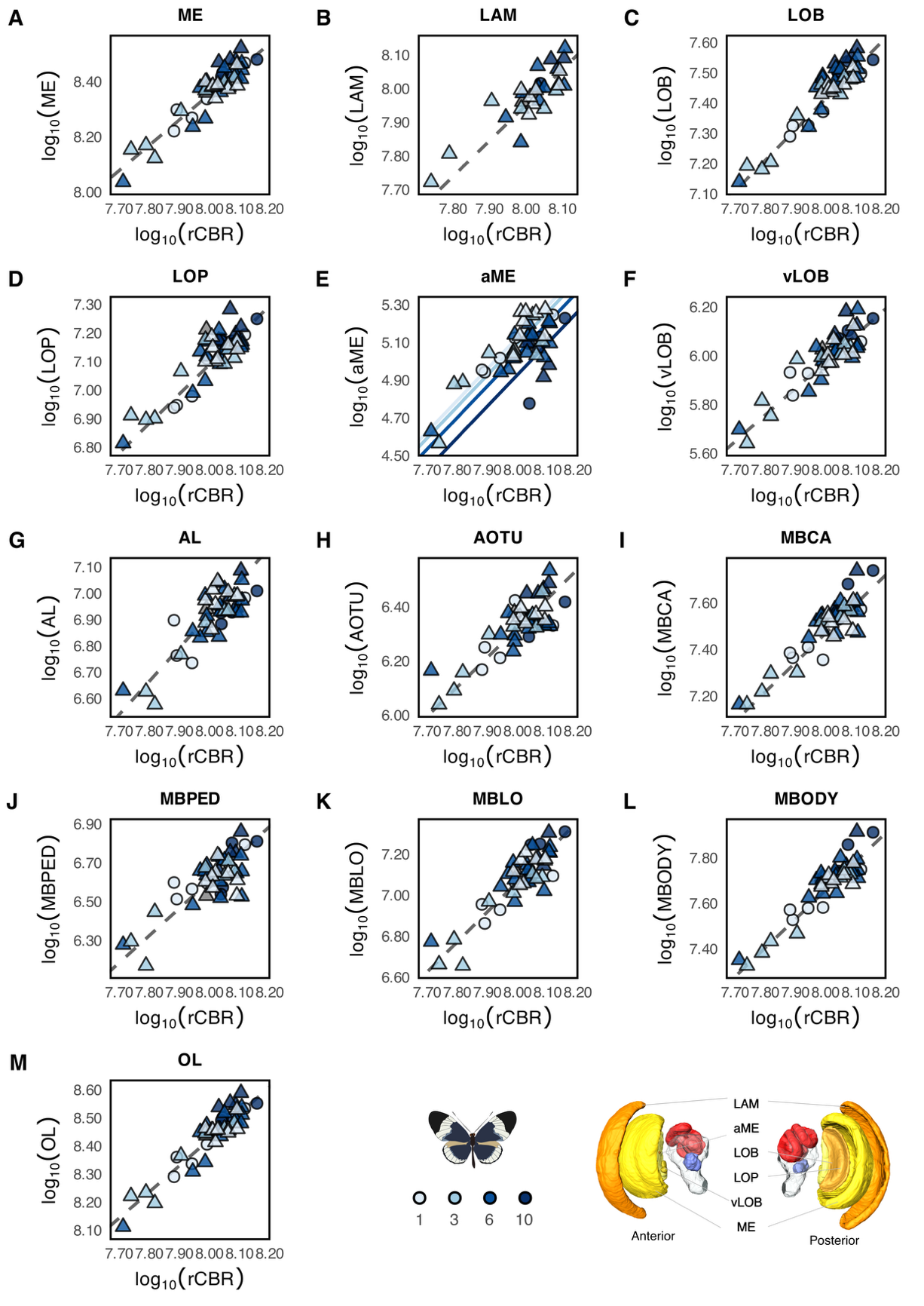

**Fig. S 7: Age-related major-axis shifts in visual and central neuropils within H. cydno.** All traits are -transformed. Scatterplots illustrate the relationships between individual neuropil volumes and the rest-of-central-brain volume (rCBR). Shapes distinguish sampling year (circles = 2022; triangles = 2023) and color saturation indicates age from 1 to 10 days (light to dark), as shown in the legend. Lines represent Standardized Major Axis (SMA) regressions; solid lines indicate elevation shifts, while dashed lines indicate a common slope and elevation.

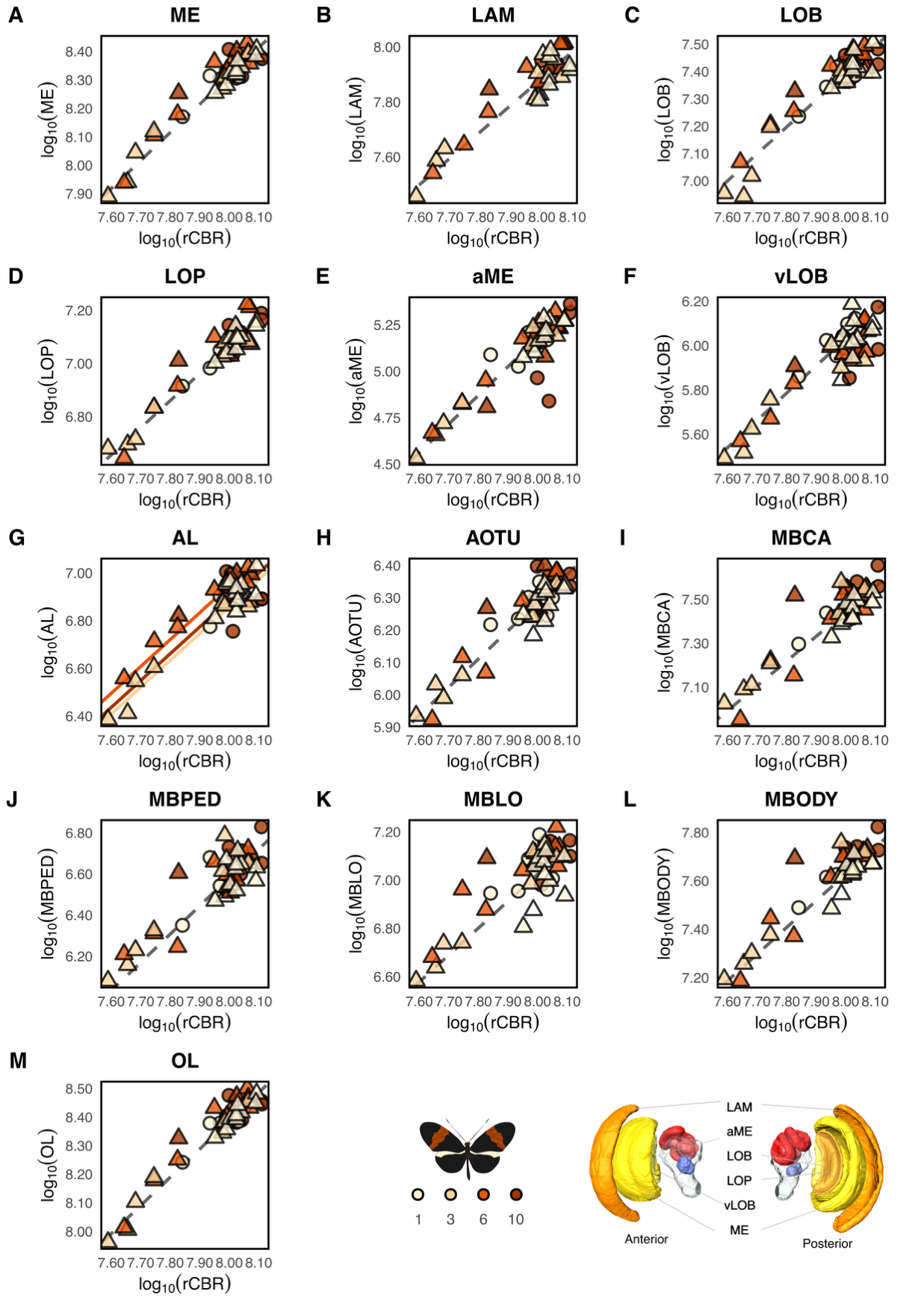

**Fig. S 8: Age-related major-axis shifts in visual and central neuropils within H. melpomene.** All traits are -transformed. Scatterplots illustrate the relationships between individual neuropil volumes and the rest-of-central-brain volume (rCBR). Shapes distinguish sampling year (circles = 2022; triangles = 2023) and color saturation indicates age from 1 to 10 days (light to dark), as shown in the legend. Lines represent Standardized Major Axis (SMA) regressions; solid lines indicate elevation shifts, while dashed lines indicate a common slope and elevation.

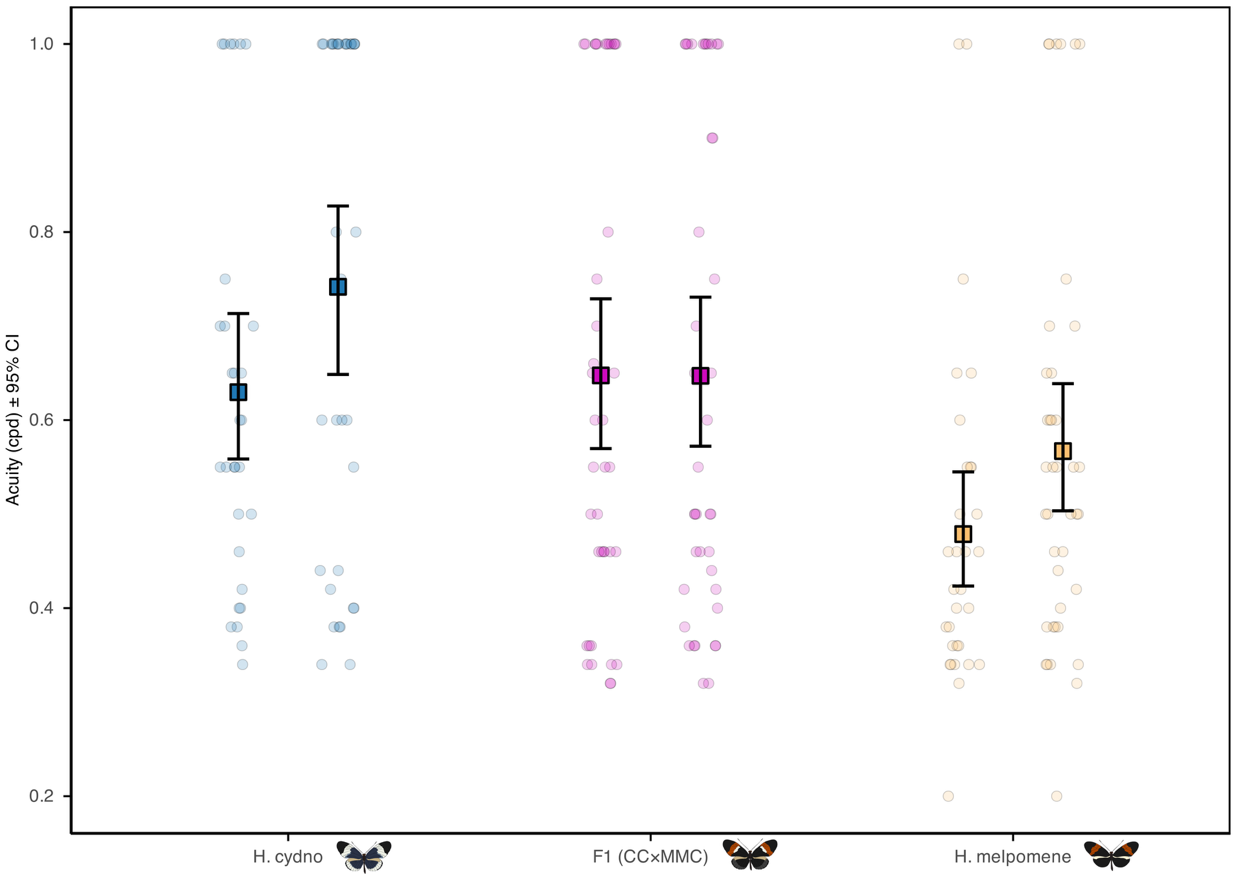

**Fig. S 9: Comparison of visual acuity in adult H. cydno, F1 hybrids and H. melpomene.** Visual acuity in females (left) and males (right) at 10 days post-eclosion. H. cydno (blue), F1 CCxMMC hybrids (magenta) H. melpomene. Points represent individuals; large squares with error bars indicate mean 95% CI. (orange).

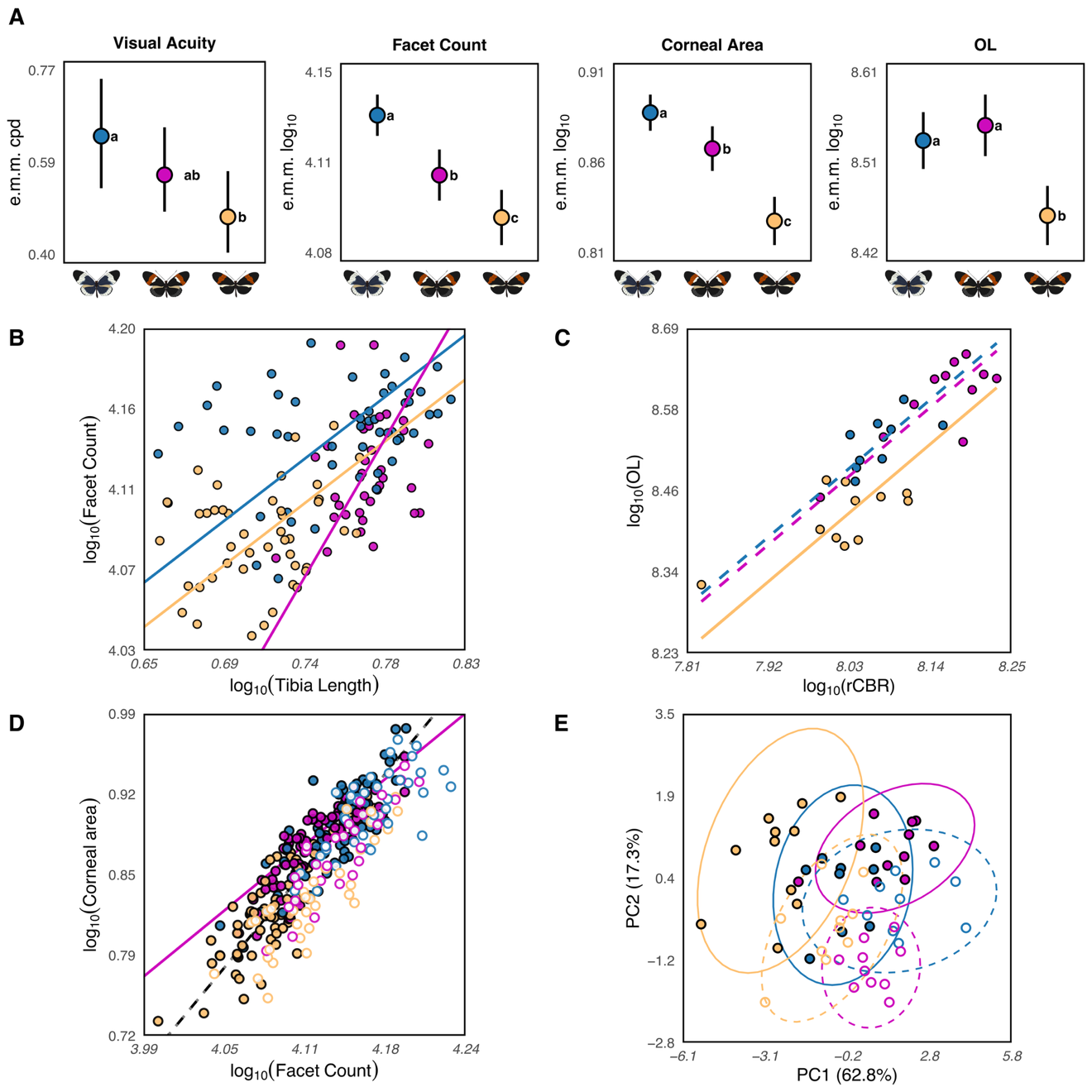

**Fig. S 10: Comparison of visual and neural traits across H. cydno, F1 hybrids and H. melpomene, incorporating data from multiple populations. scaling (A)** Estimated marginal means (e.m.m.) for visual acuity (cpd), facet count, corneal area, and optic lobe (OL) volume in H. cydno (blue),F1 hybrids (magenta), and H. melpomene (orange). Hybrids show intermediate visual acuity and corneal area, but reduced facet number relative to H. cydno and a cydno-like pattern of optic lobe investment. Letters indicate post hoc group differences (**B**) Allometric relationship between facet count and tibia length. Hybridization disrupts the conserved parental scaling, resulting in significant slope differences. (**C**) Scaling of optic lobe volume against rest-of-central-brain volume (rCBR) shows conserved slopes across groups, with differences expressed primarily as elevation shifts and hybrids resembling H. cydno. (**D-E)** Filled circles represent data from the current study (Colombia); open circles represent published data from Panama populations (Wright et al. 2024 for eyes; Montgomery et al. 2021 for brains. (**D**) Relationship between corneal area and facet count. The structural coupling observed in parental species (dashed line) is decoupled in F1 hybrids (magenta line). (**E**) Principal component analysis (PCA) of visual traits. PC1 (76.9% variance) captures the primary axis of divergence, with hybrids showing closer clustering with H. cydno in brain traits.

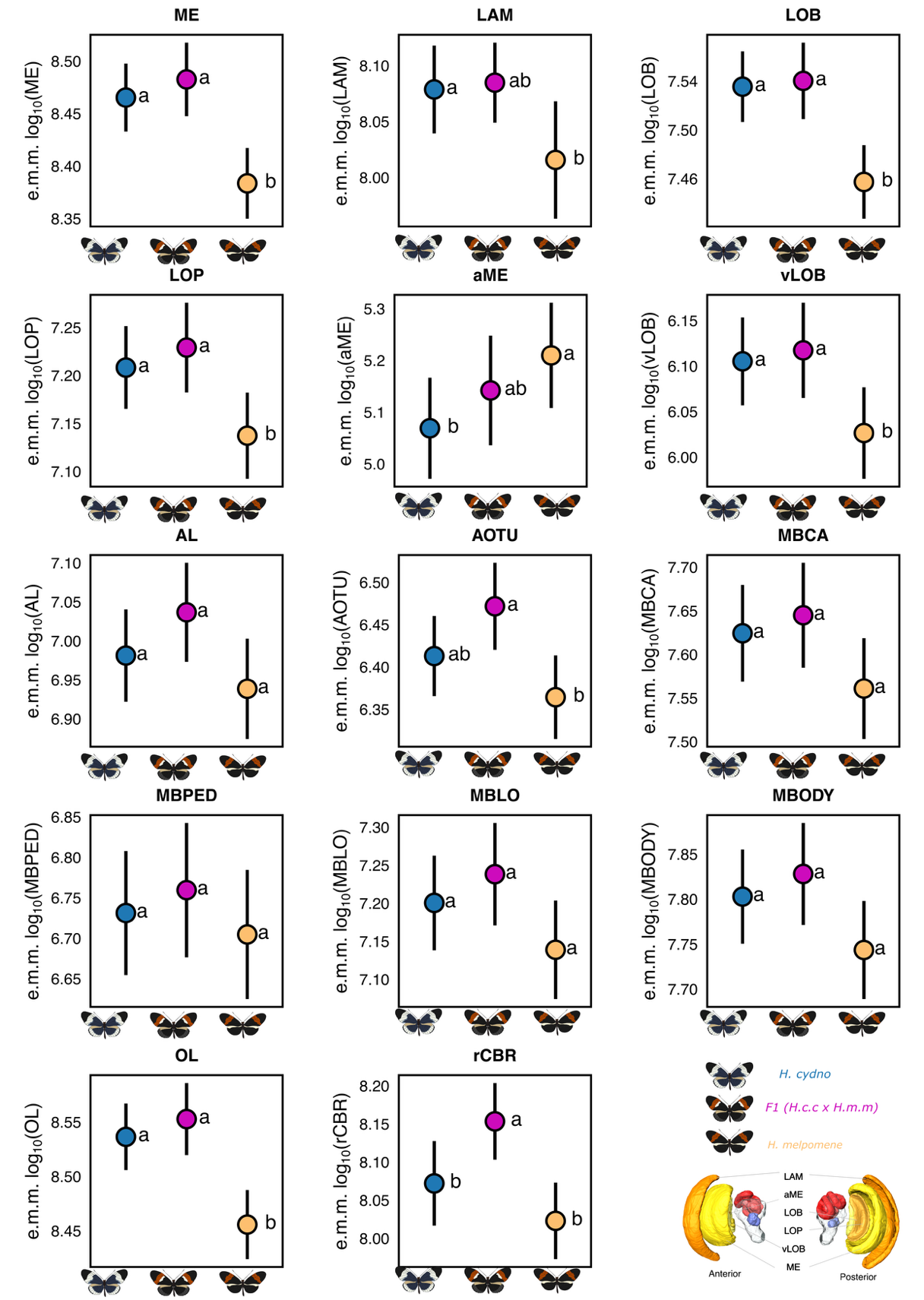

**Fig. S 11: Estimated Marginal Means (EMMs) for brain neuropil volumes across H. cydno, F1 hybrids and H. melpomene. (**A–M) All volumes are log10-transformed. Points represent the EMMs 95% credible intervals derived from Linear Mixed Models (LMMs). EMMs are corrected for sex and rest-of-central-brain volume (rCBR), representing the relative size of each neuropil independent of total brain size. Letters indicate significant group differences based on post-hoc comparisons. 3D brain reconstructions illustrate the relative positions of these regions.

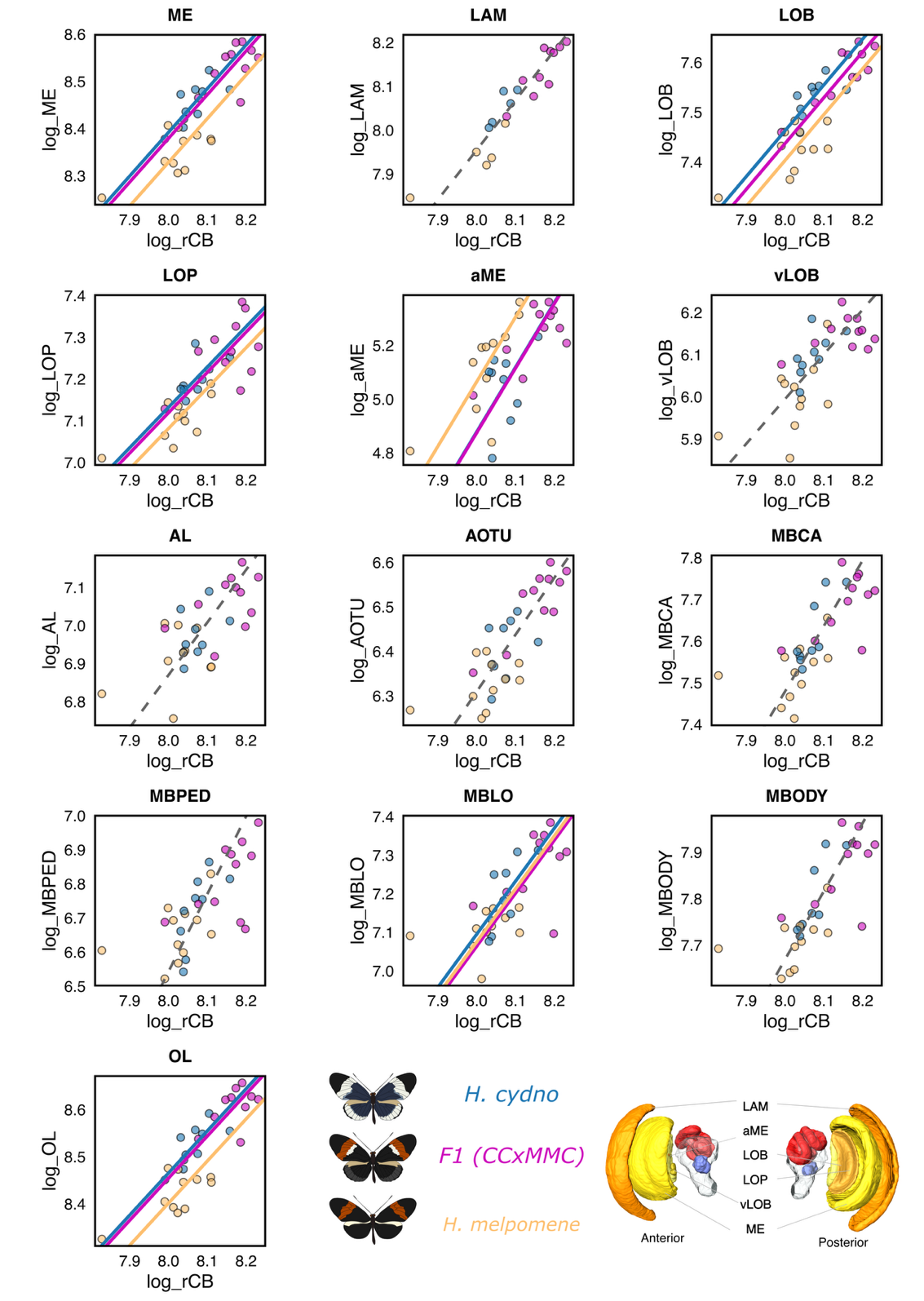

**Fig. S 12: Allometric scaling of brain neuropils across H. cydno, F1 hybrids, and H. melpomene. (A–M)** All traits are log10-transformed. Scatterplots show the relationships between individual neuropil volumes and the rest-of-central-brain volume (rCBR). Lines indicate Standardized Major Axis (SMA) regressions; solid lines indicate significant elevation shifts between groups, while dashed lines indicate a common slope and elevation (e.g., LAM, vLOB, AL). In primary visual neuropils such as the medulla (ME), lobula (LOB), and the total optic lobe (OL), F1 hybrids exhibit an upward elevation shift, scaling along a similar allometric grade as H. cydno. In contrast, the accessory medulla (aME) shows a distinct scaling relationship in hybrids compared to both parental species. 3D brain reconstructions illustrate neuropil positioning.

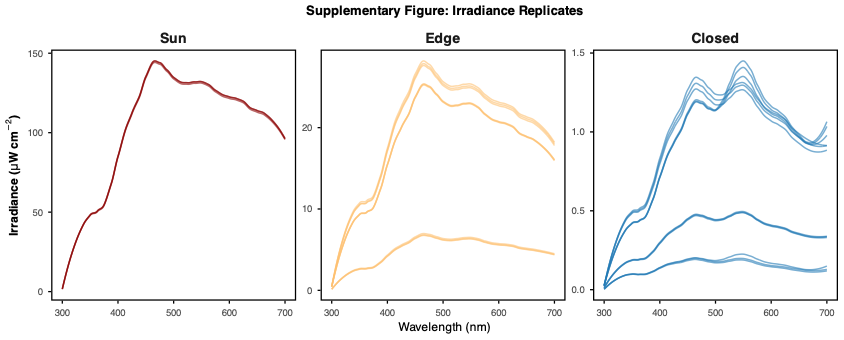

**Fig. S 13: Irradiance measurements of light environments.** Spectral irradiance (300–700 nm) recorded under (**A**) sun-exposed, (**B**) forest-edge, and (**C**) closed-canopy conditions. Each line represents a replicate measurement, showing consistent variation in both irradiance intensity and spectral distribution across habitats

| **effect** | **component** | **term** | **estimate** | **std.error** | **conf.low** | **conf.high** |
| --- | --- | --- | --- | --- | --- | --- |
| fixed | cond | (Intercept) | -0.347 | 0.093 | -0.506 | -0.151 |
| fixed | cond | Species.L | -0.080 | 0.014 | -0.108 | -0.052 |
| fixed | cond | Age | 0.012 | 0.002 | 0.008 | 0.016 |
| fixed | cond | SexMale | 0.051 | 0.022 | 0.009 | 0.093 |

**Table S 1:** Posterior estimates from a Bayesian censored Generalized Linear Model (GLM) comparing visual acuity between H. cydno (n = 169) and H. melpomene (n = 176). The model accounts for the interaction between species and age, with sex as a fixed effect and observer ID as a random effect. Estimates are reported with standard errors and 95% credible intervals. Data for acuity were log10-transformed and censored to account for detection limits. Model formula: Acuity_log | cens(cens) ~ Species * Age + Sex + (1 || Observer_live) + (1 || ID)

| **Species** | **log10_acuity** | **log10_low** | **log10_high** | **acuity_cpd** | **acuity_cpd_low** | **acuity_cpd_high** |
| --- | --- | --- | --- | --- | --- | --- |
| CC | -0.206 | -0.261 | -0.139 | 0.622 | 0.548 | 0.727 |
| MMC | -0.319 | -0.371 | -0.252 | 0.480 | 0.426 | 0.560 |

**Table S 2:** Estimated Marginal Means (EMMs) for visual acuity in H. cydno (CC) and H. melpomene (MMC). The table provides both -transformed estimates (with 95% credible intervals) and back-transformed values expressed in cycles per degree (cpd). EMMs were calculated from the Bayesian censored GLM, averaging across levels of sex and age. Sample sizes: H. cydno (n = 169), H. melpomene (n = 172).

| **effect** | **component** | **term** | **estimate** | **std.error** | **conf.low** | **conf.high** |
| --- | --- | --- | --- | --- | --- | --- |
| fixed | cond | (Intercept) | -0.342 | 0.153 | -0.648 | -0.043 |
| fixed | cond | Age | 0.016 | 0.003 | 0.010 | 0.022 |
| fixed | cond | SexMale | 0.085 | 0.036 | 0.016 | 0.156 |

**Table S 3:** Posterior estimates from a Bayesian censored Generalized Linear Model (GLM) examining the effects of age and sex on visual acuity within H. cydno (n = 169; females = 66, males = 103). The model includes age and sex as fixed effects, with observer ID and individual ID as random effects. Estimates are reported with standard errors and 95% credible intervals. Data for acuity were -transformed and censored to account for detection limits. Model formula: Acuity_log | cens(cens) ~ Age + Sex + (1 || Observer_live) + (1 || ID)

| **Sex** | **log10_acuity** | **log10_low** | **log10_high** | **acuity_cpd** | **acuity_cpd_low** | **acuity_cpd_high** |
| --- | --- | --- | --- | --- | --- | --- |
| Female | -0.2612 | -0.2985 | -0.2247 | 0.548 | 0.5029 | 0.5961 |
| Male | -0.1763 | -0.2084 | -0.1409 | 0.6664 | 0.6189 | 0.723 |

**Table S 4:** Estimated Marginal Means (EMMs) for visual acuity in H. cydno females (n = 66) and males (n = 103). The table provides both -transformed estimates (with 95% credible intervals) and back-transformed values expressed in cycles per degree (cpd). EMMs were calculated from the Bayesian censored GLM, averaging across the effect of age.

| **effect** | **component** | **term** | **estimate** | **std.error** | **conf.low** | **conf.high** |
| --- | --- | --- | --- | --- | --- | --- |
| fixed | cond | (Intercept) | -0.378 | 0.082 | -0.528 | -0.192 |
| fixed | cond | Age | 0.010 | 0.002 | 0.005 | 0.014 |
| fixed | cond | SexMale | 0.025 | 0.027 | -0.028 | 0.078 |

**Table S 5:** Posterior estimates from a Bayesian censored Generalized Linear Model (GLM) examining the effects of age and sex on visual acuity within H. melpomene (n = 172; females = 67, males = 109). The model includes age and sex as fixed effects, with observer ID and individual ID as random effects. Estimates are reported with standard errors and 95% credible intervals. Data for acuity were -transformed and censored to account for detection limits. Model formula: Acuity_log | cens(cens) ~ Age + Sex + (1 || Observer_live) + (1 || ID)

| **Sex** | **log10_acuity** | **log10_low** | **log10_high** | **acuity_cpd** | **acuity_cpd_low** | **acuity_cpd_high** |
| --- | --- | --- | --- | --- | --- | --- |
| Female | -0.329 | -0.380 | -0.265 | 0.469 | 0.417 | 0.543 |
| Male | -0.304 | -0.350 | -0.245 | 0.496 | 0.447 | 0.569 |

**Table S 6:** Estimated Marginal Means (EMMs) for visual acuity in H. melpomene females (n = 67) and males (n = 109). The table provides both -transformed estimates (with 95% credible intervals) and back-transformed values expressed in cycles per degree (cpd). EMMs were calculated from the Bayesian censored GLM, averaging across the effect of age.

| **Fixed Effect** | **Df** | **Chi-sq** | **Pr(>Chi)** | **FDR p-value** |
| --- | --- | --- | --- | --- |
| Species:Sex:Age | 1 | 0.534 | 0.387 | 0.610 |
| Species:Sex | 1 | 0.291 | 1.115 | 0.466 |
| Sex:Age | 1 | 0.115 | 2.488 | 0.229 |
| Species:Age | 3 | 0.463 | 2.569 | 0.610 |
| Age | 3 | 0.802 | 0.997 | 0.802 |
| Sex | 1 | 0.061 | 3.514 | 0.162 |
| log_Tibia_length | 1 | 0.000 | 14.198 | 0.001 |
| Species | 1 | 0.000 | 50.844 | 0.000 |

**Table S 7:** Fixed effect parameter estimates for linear model for facet count of H. cydno and H. melpomene (n= 145). For tibia length and facet count, log_10_-transofrmations were used to normalize the residuals around allometric relationships. To account for multiple tested False discovery rate (FDR) adjusted p-values are reported. Full saturated model: log_facet_count~ Species*Sex*Age+ log_Tibia_length

| **Fixed Effect** | **Df** | **Chi-sq** | **Pr(>Chi)** | **FDR p-value** |
| --- | --- | --- | --- | --- |
| Species:Sex:Age | 1 | 0.967 | 0.002 | 0.967 |
| Sex:Age | 1 | 0.346 | 0.888 | 0.461 |
| Species:Sex | 1 | 0.267 | 1.234 | 0.427 |
| Species:Age | 3 | 0.246 | 4.147 | 0.427 |
| Age | 3 | 0.229 | 4.318 | 0.427 |
| Sex | 1 | 0.407 | 0.688 | 0.465 |
| log_Tibia_length | 1 | 0.009 | 6.889 | 0.035 |
| Species | 1 | 0.000 | 50.844 | 0.000 |

**Table S 8:** Fixed effect parameter estimates for linear model for corneal area of H. cydno and H. melpomene (n= 145). For tibia length, inter-eye width and corneal area, log_10_-transofrmations were used to normalize the residuals around allometric relationships. To account for multiple tested False discovery rate (FDR) adjusted p-values are reported. Full saturated model: log_corneal_area~ Species*Sex*Age+ log_Tibia_length

|  | **Slope shift (β)** | | |  | **Elevation shift (α)** | | | | |  | **Major axis shift on common axis** | | | | |
| --- | --- | --- | --- | --- | --- | --- | --- | --- | --- | --- | --- | --- | --- | --- | --- |
|  | **LR** | **p** | **FDR** |  | **Wald-statistic** | **p** | **FDR** | **r** | **DI** |  | **Wald-statistic** | **p** | **FDR** | **r** | **DI** |
| CountTibia | 2.471 | 0.116 | 0.464 |  | 8.669 | 0.003 | 0.016 | 0.947 | CC |  | - | - | - | - | - |
| Area_Tibia | 4.887 | 0.027 | 0.135 |  | 7.360 | 0.007 | 0.027 | 0.938 | CC |  | - | - | - | - | - |
| Area_Count | 0.030 | 0.863 | 1.000 |  | 0.318 | 0.573 | 0.573 | - | - |  | 94.395 | 0.000 | 0.000 | 0.995 | CC |

**Table S 9**: Scaling relationships between eye morphology (facet count & corneal area) and body size (tibia length) for male H. cyndo and H. melpomene (n = 105). SMATR results include slope shifts (β), elevation shifts (α), and major axis shifts. FDR-corrected p-values, effect sizes (r), and direction of increase (DI) are reported. Values are for tests for grade shifts (α) and shifts along the common axis are only appropriate with a common slope.

|  | **Slope shift (β)** | | |  | **Elevation shift (α)** | | | | |  | **Major axis shift on common axis** | | | | |
| --- | --- | --- | --- | --- | --- | --- | --- | --- | --- | --- | --- | --- | --- | --- | --- |
|  | **LR** | **p** | **FDR** |  | **Wald-statistic** | **p** | **FDR** | **r** | **DI** |  | **Wald-statistic** | **p** | **FDR** | **r** | **DI** |
| CountTibia | 5.222 | 0.022 | 0.178 |  | 0.230 | 0.632 | 1.000 | - | - |  | 21.445 | 0.000 | 0.000 | 0.977 | CC |
| Area_Tibia | 5.484 | 0.019 | 0.173 |  | 2.764 | 0.096 | 0.675 | - | - |  | 34.778 | 0.000 | 0.000 | 0.986 | CC |
| Area_Count | 0.188 | 0.664 | 1.000 |  | 6.316 | 0.012 | 0.096 | 0.929 | - |  | 27.788 | 0.000 | 0.000 | 0.982 | CC |

**Table S 10**: Scaling relationships between eye morphology (facet count & corneal area) and body size (tibia length) for female H. cyndo and H. melpomene (n = 40). SMATR results include slope shifts (β), elevation shifts (α), and major axis shifts. FDR-corrected p-values, effect sizes (r), and direction of increase (DI) are reported. Values are for tests for grade shifts (α) and shifts along the common axis are only appropriate with a common slope.

| **Neuropil** | **Predictor** | **LogLh_without_factor** | **LogLh_with_factor** | **CHISQ(1)** | **p** | **FDR-p** |
| --- | --- | --- | --- | --- | --- | --- |
| ME | species | 159.397 | 169.610 | 20.427 | 0.000 | 0.000 |
| ME | age | 167.072 | 169.610 | 5.077 | 0.025 | 0.033 |
| ME | rCBR | 69.986 | 169.610 | 199.248 | < 2e-16 | < 2e-16 |
| LAM | species | 87.246 | 100.435 | 26.379 | 0.000 | 0.000 |
| LAM | age | 97.264 | 100.435 | 6.342 | 0.012 | 0.018 |
| LAM | rCBR | 45.155 | 100.435 | 110.561 | < 2e-16 | < 2e-16 |
| LOB | species | 153.850 | 162.083 | 16.466 | 0.000 | 0.000 |
| LOB | age | 158.680 | 162.083 | 6.806 | 0.009 | 0.015 |
| LOB | rCBR | 66.663 | 162.083 | 190.839 | < 2e-16 | < 2e-16 |
| LOP | species | 146.424 | 153.138 | 13.428 | 0.000 | 0.000 |
| LOP | age | 151.008 | 153.138 | 4.261 | 0.041 | 0.052 |
| LOP | rCBR | 65.705 | 153.138 | 174.867 | < 2e-16 | < 2e-16 |
| aME | species | 83.081 | 95.743 | 25.323 | 0.000 | 0.000 |
| aME | age | 84.461 | 95.743 | 22.563 | 0.000 | 0.000 |
| aME | rCBR | 28.917 | 95.743 | 133.652 | < 2e-16 | < 2e-16 |
| vLOB | species | 121.542 | 121.548 | 0.012 | 0.913 | 0.937 |
| vLOB | age | 121.542 | 121.677 | 0.271 | 0.609 | 0.658 |
| vLOB | rCBR | 48.628 | 121.542 | 145.826 | < 2e-16 | < 2e-16 |
| AL | species | 113.695 | 113.819 | 0.248 | 0.624 | 0.658 |
| AL | age | 113.695 | 113.698 | 0.005 | 0.944 | 0.944 |
| AL | rCBR | 50.593 | 113.695 | 126.206 | < 2e-16 | < 2e-16 |
| AOTU | species | 131.571 | 137.387 | 11.634 | 0.001 | 0.001 |
| AOTU | age | 137.387 | 138.475 | 2.176 | 0.147 | 0.174 |
| AOTU | rCBR | 72.342 | 137.387 | 130.091 | < 2e-16 | < 2e-16 |
| MBCA | species | 126.747 | 129.976 | 6.459 | 0.011 | 0.017 |
| MBCA | age | 124.623 | 129.976 | 10.706 | 0.001 | 0.002 |
| MBCA | rCBR | 52.844 | 129.976 | 154.266 | < 2e-16 | < 2e-16 |
| MBPED | species | 96.568 | 96.726 | 0.316 | 0.580 | 0.647 |
| MBPED | age | 96.568 | 97.720 | 2.304 | 0.133 | 0.162 |
| MBPED | rCBR | 38.656 | 96.568 | 115.826 | < 2e-16 | < 2e-16 |
| MBLO | species | 102.213 | 103.041 | 1.655 | 0.206 | 0.237 |
| MBLO | age | 98.105 | 102.213 | 8.215 | 0.004 | 0.006 |
| MBLO | rCBR | 46.180 | 102.213 | 112.065 | < 2e-16 | < 2e-16 |
| MBODY | species | 137.392 | 140.117 | 5.450 | 0.021 | 0.028 |
| MBODY | age | 133.326 | 140.117 | 13.580 | 0.000 | 0.000 |
| MBODY | rCBR | 54.685 | 140.117 | 170.864 | < 2e-16 | < 2e-16 |
| OL | species | 162.309 | 172.983 | 21.350 | 0.000 | 0.000 |
| OL | age | 170.138 | 172.983 | 5.691 | 0.018 | 0.025 |
| OL | rCBR | 69.934 | 172.983 | 206.099 | < 2e-16 | < 2e-16 |

**Table S 11**: Likelihood-ratio tests for the effects of Species, Age and rest of central brain volume (rCBR) on neuropil volumes in H. cydno and H. melpomene (n= 92). Full saturated model: log_neuropil ~ log_rCBR+ Species + Age

|  | **Slope shift (β)** | | |  | **Elevation shift (α)** | | | | |  | **Major axis shift on common axis** | | | | |
| --- | --- | --- | --- | --- | --- | --- | --- | --- | --- | --- | --- | --- | --- | --- | --- |
|  | **LR** | **p** | **FDR** |  | **Wald-statistic** | **p** | **FDR** | **r** | **DI** |  | **Wald-statistic** | **p** | **FDR** | **r** | **DI** |
| ME | 1.412 | 0.235 | 0.763 |  | 24.602 | 0.000 | 0.000 | 0.980 | CC |  | - | - | - | - | - |
| LAM | 0.165 | 0.684 | 0.868 |  | 17.573 | 0.000 | 0.000 | 0.973 | CC |  | - | - | - | - | - |
| LOB | 1.813 | 0.178 | 0.763 |  | 12.082 | 0.001 | 0.001 | 0.961 | CC |  | - | - | - | - | - |
| LOP | 0.232 | 0.630 | 0.868 |  | 10.801 | 0.001 | 0.002 | 0.957 | CC |  | - | - | - | - | - |
| aME | 0.016 | 0.901 | 0.901 |  | 33.155 | 0.000 | 0.000 | 0.985 | MMC |  | - | - | - | - | - |
| vLOB | 0.210 | 0.646 | 0.868 |  | 0.472 | 0.492 | 0.582 | 0.566 | - |  | 3.007 | 0.083 | 0.090 | 0.866 | - |
| AL | 0.585 | 0.444 | 0.840 |  | 0.010 | 0.921 | 0.921 | 0.099 | - |  | 4.107 | 0.043 | 0.050 | 0.897 | CC |
| AOTU | 0.880 | 0.348 | 0.840 |  | 8.122 | 0.004 | 0.008 | 0.944 | CC |  | - | - | - | - | - |
| MBCA | 0.063 | 0.802 | 0.868 |  | 7.412 | 0.006 | 0.011 | 0.939 | CC |  | - | - | - | - | - |
| MBPED | 0.088 | 0.767 | 0.868 |  | 0.109 | 0.741 | 0.803 | 0.314 | CC |  | 4.722 | 0.030 | 0.042 | 0.908 | CC |
| MBLO | 2.961 | 0.085 | 0.763 |  | 0.739 | 0.390 | 0.507 | 0.652 | CC |  | 4.564 | 0.033 | 0.042 | 0.906 | CC |
| MBODY | 0.565 | 0.452 | 0.840 |  | 6.075 | 0.014 | 0.020 | 0.927 | CC |  | - | - | - | - | - |
| OL | 1.552 | 0.213 | 0.763 |  | 26.175 | 0.000 | 0.000 | 0.981 | CC |  | - | - | - | - | - |

**Table S 12:** SMATR tests for species differences in neuropil allometry between H. cydno and H. melpomene (n=92) including slope shifts (β), elevation shifts (α), and major axis shifts relative to rCBR. FDR-corrected p-values, effect sizes (r), and direction of increase (DI) are reported. Values are for tests for grade shifts (α) and shifts along the common axis are only appropriate with a common slope.

|  | **Slope shift (β)** | | |  | **Elevation shift (α)** | | | | |  | **Major axis shift on common axis** | | | | |
| --- | --- | --- | --- | --- | --- | --- | --- | --- | --- | --- | --- | --- | --- | --- | --- |
|  | **LR** | **p** | **FDR** |  | **Wald-statistic** | **p** | **FDR** | **r** | **DI** |  | **Wald-statistic** | **p** | **FDR** | **r** | **DI** |
| ME | 4.249 | 0.236 | 0.497 |  | 7.601 | 0.055 | 0.179 | - | - |  | 17.510 | 0.001 | 0.002 | 0.924 | 10 |
| LAM | 3.302 | 0.347 | 0.508 |  | 3.180 | 0.365 | 0.447 | - | - |  | 8.788 | 0.032 | 0.042 | 0.863 | 10 |
| LOB | 0.362 | 0.948 | 0.948 |  | 11.630 | 0.009 | 0.057 | - | - |  | 17.516 | 0.001 | 0.002 | 0.924 | 10 |
| LOP | 3.140 | 0.370 | 0.508 |  | 5.231 | 0.156 | 0.289 | - | - |  | 16.507 | 0.001 | 0.002 | 0.920 | 10 |
| aME | 3.941 | 0.268 | 0.497 |  | 17.922 | 0.000 | 0.006 | 0.926 | 1 |  | - | - | - | - | - |
| vLOB | 0.499 | 0.919 | 0.948 |  | 6.905 | 0.075 | 0.195 | - | - |  | 19.441 | 0.000 | 0.002 | 0.931 | 10 |
| AL | 0.922 | 0.820 | 0.948 |  | 3.160 | 0.368 | 0.447 | - | - |  | 7.616 | 0.055 | 0.059 | 0.847 | - |
| AOTU | 4.414 | 0.220 | 0.497 |  | 2.051 | 0.562 | 0.609 | - | - |  | 11.044 | 0.011 | 0.017 | 0.887 | 10 |
| MBCA | 7.030 | 0.071 | 0.307 |  | 3.967 | 0.265 | 0.431 | - | - |  | 13.067 | 0.004 | 0.010 | 0.902 | 10 |
| MBPED | 10.158 | 0.017 | 0.225 |  | 0.430 | 0.934 | 0.934 | - | - |  | 8.251 | 0.041 | 0.049 | 0.856 | 10 |
| MBLO | 4.092 | 0.252 | 0.497 |  | 3.088 | 0.378 | 0.447 | - | - |  | 11.176 | 0.011 | 0.017 | 0.888 | 10 |
| MBODY | 7.269 | 0.064 | 0.307 |  | 5.301 | 0.151 | 0.289 | - | - |  | 12.591 | 0.006 | 0.010 | 0.899 | 10 |
| OL | 3.004 | 0.391 | 0.508 |  | 7.821 | 0.050 | 0.179 | - | - |  | 16.769 | 0.001 | 0.002 | 0.921 | 10 |

**Table S 13:** SMATR tests for age differences in neuropil allometry in H. cydno (n = 46) including slope shifts (β), elevation shifts (α), and major axis shifts relative to rCBR. FDR-corrected p-values, effect sizes (r), and direction of increase (DI) are reported. Values are for tests for grade shifts (α) and shifts along the common axis are only appropriate with a common slope.

|  | **Slope shift (β)** | | |  | **Elevation shift (α)** | | | | |  | **Major axis shift on common axis** | | | | |
| --- | --- | --- | --- | --- | --- | --- | --- | --- | --- | --- | --- | --- | --- | --- | --- |
|  | **LR** | **p** | **FDR** |  | **Wald-statistic** | **p** | **FDR** | **r** | **DI** |  | **Wald-statistic** | **p** | **FDR** | **r** | **DI** |
| ME | 1.021 | 0.796 | 0.876 |  | 5.176 | 0.159 | 0.286 | - | - |  | 8.952 | 0.030 | 0.038 | 0.948 | 10 |
| LAM | 5.190 | 0.158 | 0.761 |  | 3.431 | 0.330 | 0.390 | - | - |  | 7.275 | 0.064 | 0.064 | - | - |
| LOB | 2.841 | 0.417 | 0.761 |  | 4.611 | 0.203 | 0.286 | - | - |  | 9.079 | 0.028 | 0.038 | 0.949 | 10 |
| LOP | 2.539 | 0.468 | 0.761 |  | 1.357 | 0.716 | 0.775 | - | - |  | 8.782 | 0.032 | 0.038 | 0.948 | 10 |
| aME | 5.283 | 0.152 | 0.761 |  | 8.489 | 0.037 | 0.240 | - | - |  | 8.319 | 0.040 | 0.043 | 0.945 | 1 |
| vLOB | 0.887 | 0.829 | 0.876 |  | 5.702 | 0.127 | 0.286 | - | - |  | 10.986 | 0.012 | 0.026 | 0.957 | 1 |
| AL | 2.019 | 0.568 | 0.821 |  | 13.742 | 0.003 | 0.043 | 0.906 | 6 |  | - | - | - | - | - |
| AOTU | 4.741 | 0.192 | 0.761 |  | 0.797 | 0.850 | 0.850 | - | - |  | 10.940 | 0.012 | 0.026 | 0.957 | 10 |
| MBCA | 3.172 | 0.366 | 0.761 |  | 6.443 | 0.092 | 0.286 | - | - |  | 12.179 | 0.007 | 0.022 | 0.961 | 10 |
| MBPED | 0.688 | 0.876 | 0.876 |  | 4.418 | 0.220 | 0.286 | - | - |  | 14.271 | 0.003 | 0.017 | 0.967 | 10 |
| MBLO | 3.658 | 0.301 | 0.761 |  | 4.815 | 0.186 | 0.286 | - | - |  | 15.130 | 0.002 | 0.017 | 0.969 | 10 |
| MBODY | 3.236 | 0.357 | 0.761 |  | 5.102 | 0.165 | 0.286 | - | - |  | 12.950 | 0.005 | 0.021 | 0.963 | 10 |
| OL | 1.426 | 0.700 | 0.876 |  | 4.861 | 0.182 | 0.286 | - | - |  | 9.302 | 0.026 | 0.038 | 0.950 | 10 |

**Table S 14:** SMATR tests for age differences differnces in neuropil allometry in H. melpomene (n = 46) including slope shifts (β), elevation shifts (α), and major axis shifts relative to rCBR. FDR-corrected p-values, effect sizes (r), and direction of increase (DI) are reported. Values are for tests for grade shifts (α) and shifts along the common axis are only appropriate with a common slope.

| **effect** | **component** | **term** | **estimate** | **std.error** | **conf.low** | **conf.high** |
| --- | --- | --- | --- | --- | --- | --- |
| fixed | cond | (Intercept) | -0.286 | 0.136 | -0.572 | -0.018 |
| fixed | cond | Species.L | -0.087 | 0.026 | -0.137 | -0.036 |
| fixed | cond | Species.Q | -0.006 | 0.026 | -0.058 | 0.047 |
| fixed | cond | SexMale | 0.066 | 0.029 | 0.009 | 0.124 |

**Table S 15:** Posterior estimates from a Bayesian censored Generalized Linear Model (GLM) comparing visual acuity across 10-day old H. cydno(n = 64), F1 CCxMMC hybrids (n = 81), and H. melpomene (n = 73). The model includes species, and sex as fixed effects, with observer ID as a random effect. Estimates are reported with standard errors and 95% credible intervals. Data for acuity were -transformed and censored to account for detection limits. Model formula: Acuity_log | cens(cens) ~ Species + Sex + (1 || Observer_live)

| **Species** | **log10_acuity** | **log10_low** | **log10_high** | **acuity_cpd** | **acuity_cpd_low** | **acuity_cpd_high** |
| --- | --- | --- | --- | --- | --- | --- |
| CC | -0.196 | -0.272 | -0.125 | 0.638 | 0.535 | 0.750 |
| CCxMMC | -0.251 | -0.311 | -0.184 | 0.561 | 0.489 | 0.655 |
| MMC | -0.320 | -0.389 | -0.245 | 0.479 | 0.408 | 0.569 |

**Table S 16:** Estimated Marginal Means (EMMs) for visual acuity in H. cydno (CC; n = 64), F1 CCxMMC hybrids (n = 81), and H. melpomene (MMC; n = 73). The table provides model-predicted means and 95% credible intervals on both the scale and back-transformed into cycles per degree (cpd). EMMs were calculated from the Bayesian censored GLM, averaging across the effects of sex.

| **Fixed Effect** | **Df** | **Chi-sq** | **Pr(>Chi)** | **FDR p-value** |
| --- | --- | --- | --- | --- |
| Species:Sex | 2 | 0.119 | 4.252 | 0.119 |
| Sex | 1 | 0.014 | 6.047 | 0.019 |
| log_Tibia_length | 1 | 0.000 | 20.483 | 0.000 |
| Species | 2 | 0.000 | 55.262 | 0.000 |

**Table S 17:** Fixed effect parameter estimates for linear model for facet count of H. cydno, F1 hybrids (CCxMMC) and H. melpomene (n= 205). For tibia length and facet count, log_10_-transofrmations were used to normalize the residuals around allometric relationships. To account for multiple tested False discovery rate (FDR) adjusted p-values are reported. Full saturated model: log_facet_count~ Species*Sex* + log_Tibia_length

| **Comparison** | **estimate** | **SE** | **df** | **t-ratio** | **p-value** |
| --- | --- | --- | --- | --- | --- |
| CC - CCxMMC | 0.023 | 0.005 | 185 | 4.480 | 0.000 |
| CC - MMC | 0.040 | 0.005 | 185 | 7.317 | 0.000 |
| CCxMMC - MMC | 0.016 | 0.007 | 185 | 2.464 | 0.044 |

**Table S 18:** Bonferroni-corrected pairwise comparisons for the linear model of facet count in H. cydno, F1 hybrids (CCxMMC), and H. melpomene (n = 205). To normalize residuals and account for allometric relationships, both tibia length and facet count were -transformed. Reported p-values are adjusted for multiple comparisons. Full model: log_facet_count~ Species*Sex + log_Tibia_length

| **Fixed Effect** | **Df** | **Chi-sq** | **Pr(>Chi)** | **FDR p-value** |
| --- | --- | --- | --- | --- |
| log_Tibia_length | 1 | 0.000 | 20.483 | 0.000 |
| Species | 2 | 0.000 | 55.262 | 0.000 |

**Table S 19:** Fixed effect parameter estimates for linear model for facet count of male H. cydno, F1 hybrids (CCxMMC) and H. melpomene (n= 142). For tibia length and facet count, log_10_-transofrmations were used to normalize the residuals around allometric relationships. To account for multiple tested False discovery rate (FDR) adjusted p-values are reported. Full saturated model: log_facet_count~ Species+ log_Tibia_length

| **Comparison** | **estimate** | **SE** | **df** | **t-ratio** | **p-value** |
| --- | --- | --- | --- | --- | --- |
| CC - CCxMMC | 0.025 | 0.005 | 186.000 | 4.862 | 0.000 |
| CC - MMC | 0.038 | 0.005 | 186.000 | 6.929 | 0.000 |
| CCxMMC - MMC | 0.012 | 0.007 | 186.000 | 1.889 | 0.181 |

**Table S 20:** Bonferroni-corrected pairwise comparisons for the linear model of facet count in male H. cydno, F1 hybrids (CCxMMC), and H. melpomene (n = 142). To normalize residuals and account for allometric relationships, both tibia length and corneal area were log10-transformed. Reported p-values are adjusted for multiple comparisons. Full model: log_corneal_area ~ Species + log_Tibia_length

| **Fixed Effect** | **Df** | **Chi-sq** | **Pr(>Chi)** | **FDR p-value** |
| --- | --- | --- | --- | --- |
| Species:Sex | 2 | 0.134 | 4.022 | 0.134 |
| Sex | 1 | 0.042 | 4.119 | 0.057 |
| log_Tibia_length | 1 | 0.000 | 12.821 | 0.001 |
| Species | 2 | 0.000 | 55.262 | 0.000 |

**Table S 21:** Fixed effect parameter estimates for linear model for corneal area of H. cydno, F1 hybrids (CCxMMC) and H. melpomene (n= 205). For tibia length and corneal are, log_10_-transofrmations were used to normalize the residuals around allometric relationships. To account for multiple tested False discovery rate (FDR) adjusted p-values are reported. Full saturated model: log_corneal_area ~ Species*Sex + log_Tibia_length

| **Comparison** | **estimate** | **SE** | **df** | **t-ratio** | **p-value** |
| --- | --- | --- | --- | --- | --- |
| CC - CCxMMC | 0.021 | 0.007 | 185 | 3.076 | 0.007 |
| CC - MMC | 0.062 | 0.007 | 185 | 8.859 | 0.000 |
| CCxMMC - MMC | 0.041 | 0.009 | 185 | 4.808 | 0.000 |

**Table S 22:** Bonferroni-corrected pairwise comparisons for the linear model of facet count in H. cydno, F1 hybrids (CCxMMC), and H. melpomene (n = 205). To normalize residuals and account for allometric relationships, both tibia length and corneal area were log10-transformed. Reported p-values are adjusted for multiple comparisons. Full model: log_corneal_area ~ Species*Sex + log_Tibia_length

| **Fixed Effect** | **Df** | **Chi-sq** | **Pr(>Chi)** | **FDR p-value** |
| --- | --- | --- | --- | --- |
| log_Tibia_length | 1 | 0.000 | 12.821 | 0.000 |
| Species | 2 | 0.000 | 55.262 | 0.000 |

**Table S 23:** Fixed effect parameter estimates for linear model for corneal area of male H. cydno, F1 hybrids (CCxMMC) and H. melpomene (n= 142). For tibia length and corneal are, log_10_-transofrmations were used to normalize the residuals around allometric relationships. To account for multiple tested False discovery rate (FDR) adjusted p-values are reported. Full saturated model: log_corneal_area ~ Species+ log_Tibia_length

| **Comparison** | **estimate** | **SE** | **df** | **t-ratio** | **p-value** |
| --- | --- | --- | --- | --- | --- |
| CC - CCxMMC | 0.023 | 0.007 | 186.000 | 3.408 | 0.002 |
| CC - MMC | 0.060 | 0.007 | 186.000 | 8.581 | 0.000 |
| CCxMMC - MMC | 0.037 | 0.008 | 186.000 | 4.410 | 0.000 |

**Table S 24:** Bonferroni-corrected pairwise comparisons for the linear model of facet count in male H. cydno, F1 hybrids (CCxMMC), and H. melpomene (n = 142). To normalize residuals and account for allometric relationships, both tibia length and corneal area were log10-transformed. Reported p-values are adjusted for multiple comparisons. Full model: log_corneal_area ~ Species+ log_Tibia_length

|  | **Slope shift (β)** | | |  | **Elevation shift (α)** | | | | |  | **Major axis shift on common axis** | | | | |
| --- | --- | --- | --- | --- | --- | --- | --- | --- | --- | --- | --- | --- | --- | --- | --- |
|  | **LR** | **p** | **FDR** |  | **Wald-statistic** | **p** | **FDR** | **r** | **DI** |  | **Wald-statistic** | **p** | **FDR** | **r** | **DI** |
| CountTibia | 8.078 | 0.018 | 0.018 |  | - | - | - | - | - |  | - | - | - | - | - |
| Area_Tibia | 13.433 | 0.001 | 0.002 |  | - | - | - | - | - |  | - | - | - | - | - |
| Area_Count | 15.198 | 0.001 | 0.002 |  | - | - | - | - | - |  | - | - | - | - | - |

**Table S 25**: Scaling relationships between eye morphology (facet count & corneal area) and body size (tibia length) for male H. cydno, F1 hybrids (CCxMMC), and H. melpomene (n = 142). SMATR results include slope shifts (β), elevation shifts (α), and major axis shifts. FDR-corrected p-values, effect sizes (r), and direction of increase (DI) are reported. Values are for tests for grade shifts (α) and shifts along the common axis are only appropriate with a common slope.

|  |  | **Slope shift (β)** | | |  | **Elevation shift (α)** | | | | |  | **Major axis shift on common axis** | | | | |
| --- | --- | --- | --- | --- | --- | --- | --- | --- | --- | --- | --- | --- | --- | --- | --- | --- |
| **Relationship** | **Comparison** | **LR** | **p** | **FDR** |  | **Wald-statistic** | **p** | **FDR** | **r** | **DI** |  | **Wald-statistic** | **p** | **FDR** | **r** | **DI** |
| Facet_Count ~ Tibia | CC vs MMC | 2.471 | 0.116 | 0.348 |  | 8.669 | 0.003 | 0.010 | 0.947 | CC |  | - | - | - | - | - |
| Facet_Count ~ Tibia | CC vs CCxMMC | 13.429 | 0.000 | 0.001 |  | - | - | - | - | - |  | - | - | - | - | - |
| Facet_Count ~ Tibia | MMC vs CCxMMC | 4.590 | 0.032 | 0.096 |  | - | - | - | - | - |  | - | - | - | - | - |
| Corneal_Area ~ Tibia | CC vs MMC | 4.887 | 0.027 | 0.081 |  | - | - | - | - | - |  | - | - | - | - | - |
| Corneal_Area ~ Tibia | CC vs CCxMMC | 6.594 | 0.010 | 0.031 |  | - | - | - | - | - |  | - | - | - | - | - |
| Corneal_Area ~ Tibia | MMC vs CCxMMC | 0.320 | 0.572 | 1.000 |  | 1.456 | 0.228 | 0.683 | 0.770 | MMC |  | 90.224 | 0.000 | 0.000 | 0.995 | CCxMMC |
| Area ~ Facet_Count | CC vs MMC | 0.002 | 0.967 | 1.000 |  | 2.061 | 0.151 | 0.453 | - | - |  | 96.481 | 0.000 | 0.000 | 0.995 | CC |
| Area ~ Facet_Count | CC vs CCxMMC | 12.649 | 0.000 | 0.001 |  | - | - | - | - | - |  | - | - | - | - | - |
| Area ~ Facet_Count | MMC vs CCxMMC | 12.073 | 0.001 | 0.002 |  | - | - | - | - | - |  | - | - | - | - | - |

**Table S 26**: Pairwise scaling relationships between eye morphology (facet count & corneal area) and body size (tibia length) for male H. cydno, F1 hybrids (CCxMMC), and H. melpomene (n = 142). SMATR results include slope shifts (β), elevation shifts (α), and major axis shifts. FDR-corrected p-values, effect sizes (r), and direction of increase (DI) are reported. Values are for tests for grade shifts (α) and shifts along the common axis are only appropriate with a common slope.

| **Neuropil** | **Predictor** | **LogLh_without_factor** | **LogLh_with_factor** | **CHISQ(1)** | **p** | **FDR-p** |
| --- | --- | --- | --- | --- | --- | --- |
| ME | species | 54.866 | 65.767 | 21.803 | < 0.001 | < 0.001 |
| ME | rCBR | 54.064 | 65.767 | 23.407 | < 0.001 | < 0.001 |
| LAM | species | 39.449 | 47.545 | 16.191 | < 0.001 | < 0.001 |
| LAM | rCBR | 36.752 | 47.545 | 21.585 | < 0.001 | < 0.001 |
| LOB | species | 58.543 | 69.131 | 21.175 | < 0.001 | < 0.001 |
| LOB | rCBR | 55.756 | 69.131 | 26.749 | < 0.001 | < 0.001 |
| LOP | species | 48.725 | 56.128 | 14.807 | < 0.001 | < 0.001 |
| LOP | rCBR | 49.803 | 56.128 | 12.650 | < 0.001 | < 0.001 |
| aME | species | 24.867 | 28.712 | 7.690 | 0.022 | 0.025 |
| aME | rCBR | 19.151 | 28.712 | 19.122 | < 0.001 | < 0.001 |
| vLOB | species | 46.051 | 50.023 | 7.944 | 0.019 | 0.023 |
| vLOB | rCBR | 34.629 | 46.051 | 22.843 | < 0.001 | < 0.001 |
| AL | species | 40.114 | 42.234 | 4.239 | 0.137 | 0.143 |
| AL | rCBR | 31.013 | 40.114 | 18.202 | < 0.001 | < 0.001 |
| AOTU | species | 46.644 | 51.613 | 9.939 | 0.006 | 0.008 |
| AOTU | rCBR | 44.080 | 51.613 | 15.066 | < 0.001 | < 0.001 |
| MBCA | species | 42.636 | 46.688 | 8.103 | 0.018 | 0.023 |
| MBCA | rCBR | 42.064 | 46.688 | 9.247 | 0.002 | 0.003 |
| MBPED | species | 33.409 | 34.268 | 1.718 | 0.460 | 0.460 |
| MBPED | rCBR | 24.562 | 33.409 | 17.694 | < 0.001 | < 0.001 |
| MBLO | species | 41.116 | 43.843 | 5.454 | 0.074 | 0.080 |
| MBLO | rCBR | 30.319 | 41.116 | 21.594 | < 0.001 | < 0.001 |
| MBODY | species | 45.799 | 49.804 | 8.009 | 0.019 | 0.023 |
| MBODY | rCBR | 44.111 | 49.804 | 11.385 | < 0.001 | < 0.001 |
| OL | species | 56.142 | 67.828 | 23.371 | < 0.001 | < 0.001 |
| OL | rCBR | 55.126 | 67.828 | 25.405 | < 0.001 | < 0.001 |

**Table S 27**: Likelihood-ratio tests for the effects of Species and rest of central brain volume (rCBR) on neuropil volumes in 10 day old H. cydno, F1 hybrids (CCxMMC), and H. melpomene (n= 34). Full saturated model: log_neuropil ~ log_rCBR+ Species

| **Neuropil** | **contrast** | **estimate** | **SE** | **df** | **t.-ratio** | **p-value** |
| --- | --- | --- | --- | --- | --- | --- |
| ME | CC - CCxMMC | -0.021 | 0.018 | 30.000 | -1.177 | 0.745 |
| ME | CC - MMC | 0.073 | 0.016 | 30.000 | 4.505 | 0.000 |
| ME | CCxMMC - MMC | 0.095 | 0.021 | 30.000 | 4.602 | 0.000 |
| LAM | CC - CCxMMC | -0.001 | 0.021 | 19.000 | -0.032 | 1.000 |
| LAM | CC - MMC | 0.089 | 0.021 | 19.000 | 4.369 | 0.001 |
| LAM | CCxMMC - MMC | 0.090 | 0.028 | 19.000 | 3.254 | 0.013 |
| LOB | CC - CCxMMC | -0.005 | 0.016 | 30.000 | -0.303 | 1.000 |
| LOB | CC - MMC | 0.070 | 0.015 | 30.000 | 4.780 | 0.000 |
| LOB | CCxMMC - MMC | 0.075 | 0.019 | 30.000 | 4.051 | 0.001 |
| LOP | CC - CCxMMC | -0.025 | 0.024 | 30.000 | -1.045 | 0.913 |
| LOP | CC - MMC | 0.074 | 0.022 | 30.000 | 3.443 | 0.005 |
| LOP | CCxMMC - MMC | 0.099 | 0.027 | 30.000 | 3.645 | 0.003 |
| aME | CC - CCxMMC | -0.056 | 0.054 | 30.000 | -1.041 | 0.919 |
| aME | CC - MMC | -0.133 | 0.048 | 30.000 | -2.747 | 0.030 |
| aME | CCxMMC - MMC | -0.077 | 0.061 | 30.000 | -1.258 | 0.654 |
| vLOB | CC - CCxMMC | -0.005 | 0.029 | 30.000 | -0.171 | 1.000 |
| vLOB | CC - MMC | 0.068 | 0.026 | 30.000 | 2.637 | 0.039 |
| vLOB | CCxMMC - MMC | 0.073 | 0.033 | 30.000 | 2.237 | 0.099 |
| AL | CC - CCxMMC | -0.054 | 0.035 | 29.000 | -1.540 | 0.403 |
| AL | CC - MMC | 0.025 | 0.032 | 29.000 | 0.779 | 1.000 |
| AL | CCxMMC - MMC | 0.079 | 0.040 | 29.000 | 1.961 | 0.179 |
| AOTU | CC - CCxMMC | -0.048 | 0.027 | 30.000 | -1.736 | 0.279 |
| AOTU | CC - MMC | 0.051 | 0.025 | 30.000 | 2.066 | 0.143 |
| AOTU | CCxMMC - MMC | 0.099 | 0.031 | 30.000 | 3.162 | 0.011 |
| MBCA | CC - CCxMMC | -0.034 | 0.032 | 30.000 | -1.071 | 0.878 |
| MBCA | CC - MMC | 0.063 | 0.029 | 30.000 | 2.212 | 0.104 |
| MBCA | CCxMMC - MMC | 0.097 | 0.036 | 30.000 | 2.694 | 0.034 |
| MBPED | CC - CCxMMC | -0.050 | 0.046 | 30.000 | -1.090 | 0.854 |
| MBPED | CC - MMC | 0.011 | 0.041 | 30.000 | 0.267 | 1.000 |
| MBPED | CCxMMC - MMC | 0.061 | 0.052 | 30.000 | 1.170 | 0.753 |
| MBLO | CC - CCxMMC | -0.037 | 0.034 | 30.000 | -1.079 | 0.868 |
| MBLO | CC - MMC | 0.050 | 0.031 | 30.000 | 1.618 | 0.348 |
| MBLO | CCxMMC - MMC | 0.087 | 0.039 | 30.000 | 2.230 | 0.100 |
| MBODY | CC - CCxMMC | -0.035 | 0.029 | 30.000 | -1.218 | 0.698 |
| MBODY | CC - MMC | 0.054 | 0.026 | 30.000 | 2.088 | 0.136 |
| MBODY | CCxMMC - MMC | 0.090 | 0.033 | 30.000 | 2.725 | 0.032 |
| OL | CC - CCxMMC | -0.020 | 0.017 | 30.000 | -1.168 | 0.756 |
| OL | CC - MMC | 0.073 | 0.015 | 30.000 | 4.758 | 0.000 |
| OL | CCxMMC - MMC | 0.093 | 0.019 | 30.000 | 4.794 | 0.000 |

**Table S 28**: Bonferroni-corrected pairwise comparisons for the linear model effects of Species and rest of central brain volume (rCBR) on neuropil volumes in 10-day old H. cydno, F1 hybrids (CCxMMC), and H. melpomene (n= 34).
Full saturated model: log_neuropil ~ log_rCBR+ Species + Age

|  | **Slope shift (β)** | | |  | **Elevation shift (α)** | | | | |  | **Major axis shift on common axis** | | | | |
| --- | --- | --- | --- | --- | --- | --- | --- | --- | --- | --- | --- | --- | --- | --- | --- |
|  | **LR** | **p** | **FDR** |  | **Wald-statistic** | **p** | **FDR** | **r** | **DI** |  | **Wald-statistic** | **p** | **FDR** | **r** | **DI** |
| ME | 1.2041 | 0.5477 | 0.6638 |  | 10.889 | 0.004 | 0.014 | 0.919 | CC |  | - | - | - | - | - |
| LAM | 3.8508 | 0.1458 | 0.4056 |  | 3.752 | 0.153 | 0.270 | - | - |  | 24.257 | 0.000 | 0.000 | 0.961 | CCxMMC |
| LOB | 1.0827 | 0.5820 | 0.6638 |  | 12.323 | 0.002 | 0.009 | 0.928 | CC |  | - | - | - | - | - |
| LOP | 0.9796 | 0.6128 | 0.6638 |  | 7.097 | 0.029 | 0.075 | - | - |  | 28.274 | 0.000 | 0.000 | 0.966 | CCxMMC |
| aME | 1.6126 | 0.4465 | 0.6638 |  | 13.519 | 0.001 | 0.009 | 0.933 | MMC |  | - | - | - | - | - |
| vLOB | 2.5844 | 0.2747 | 0.5101 |  | 5.584 | 0.061 | 0.133 | - | - |  | 32.757 | 0.000 | 0.000 | 0.971 | CCxMMC |
| AL | 0.8044 | 0.6689 | 0.6689 |  | 0.349 | 0.840 | 0.840 | - | - |  | 21.813 | 0.000 | 0.000 | 0.957 | CCxMMC |
| AOTU | 2.7733 | 0.2499 | 0.5101 |  | 0.652 | 0.722 | 0.782 | - | - |  | 33.464 | 0.000 | 0.000 | 0.971 | CCxMMC |
| MBCA | 3.7156 | 0.1560 | 0.4056 |  | 3.587 | 0.166 | 0.270 | - | - |  | 25.937 | 0.000 | 0.000 | 0.964 | CCxMMC |
| MBPED | 4.1001 | 0.1287 | 0.4056 |  | 3.050 | 0.218 | 0.314 | - | - |  | 18.901 | 0.000 | 0.000 | 0.951 | CCxMMC |
| MBLO | 6.2763 | 0.0434 | 0.4056 |  | 0.765 | 0.682 | 0.782 | - | - |  | 27.419 | 0.000 | 0.000 | 0.965 | CCxMMC |
| MBODY | 4.0098 | 0.1347 | 0.4056 |  | 2.705 | 0.259 | 0.336 | - | - |  | 26.970 | 0.000 | 0.000 | 0.965 | CCxMMC |
| OL | 1.3565 | 0.5075 | 0.6638 |  | 12.423 | 0.002 | 0.009 | 0.928 | CC |  | - | - | - | - | - |

**Table S 29:** SMATR tests for species differences in neuropil allometry in 10-day old H. cydno, F1 hybrids (CCxMMC), and H. melpomene (n= 34), including slope shifts (β), elevation shifts (α), and major axis shifts relative to rCBR. FDR-corrected p-values, effect sizes (r), and direction of increase (DI) are reported. Values are for tests for grade shifts (α) and shifts along the common axis are only appropriate with a common slope.

|  |  | **Slope shift (β)** | | |  | **Elevation shift (α)** | | | | |  | **Major axis shift on common axis** | | | | |
| --- | --- | --- | --- | --- | --- | --- | --- | --- | --- | --- | --- | --- | --- | --- | --- | --- |
|  |  | **LR** | **p** | **FDR** |  | **Wald-statistic** | **p** | **FDR** | **r** | **DI** |  | **Wald-statistic** | **p** | **FDR** | **r** | **DI** |
| CC vs MMC | ME | 1.143 | 0.285 | 0.855 |  | 9.679 | 0.002 | 0.006 | 0.952 | CC |  | - | - | - | - | - |
| CC vs CCxMMC | ME | 0.631 | 0.427 | 1.000 |  | 0.627 | 0.428 | 1.000 | - | - |  | 13.408 | 0.000 | 0.001 | 0.965 | CCxMMC |
| MMC vs CCxMMC | ME | 0.169 | 0.681 | 1.000 |  | 6.256 | 0.012 | 0.037 | 0.929 | CCxMMC |  | - | - | - | - | - |
| CC vs MMC | LAM | 3.276 | 0.070 | 0.211 |  | 4.353 | 0.037 | 0.111 | - | - |  | 5.014 | 0.025 | 0.075 | - | - |
| CC vs CCxMMC | LAM | 0.005 | 0.945 | 1.000 |  | 3.877 | 0.049 | 0.147 | - | - |  | 15.079 | 0.000 | 0.000 | 0.968 | CCxMMC |
| MMC vs CCxMMC | LAM | 3.719 | 0.054 | 0.161 |  | 0.275 | 0.600 | 1.000 | - | - |  | 17.132 | 0.000 | 0.000 | 0.972 | CCxMMC |
| CC vs MMC | LOB | 0.974 | 0.324 | 0.971 |  | 8.129 | 0.004 | 0.013 | 0.944 | CC |  | - | - | - | - | - |
| CC vs CCxMMC | LOB | 0.943 | 0.332 | 0.995 |  | 2.647 | 0.104 | 0.311 | - | - |  | 9.742 | 0.002 | 0.005 | 0.952 | CCxMMC |
| MMC vs CCxMMC | LOB | 0.035 | 0.852 | 1.000 |  | 3.736 | 0.053 | 0.160 | - | - |  | 32.957 | 0.000 | 0.000 | 0.985 | CCxMMC |
| CC vs MMC | LOP | 0.035 | 0.851 | 1.000 |  | 8.297 | 0.004 | 0.012 | 0.945 | CC |  | - | - | - | - | - |
| CC vs CCxMMC | LOP | 0.905 | 0.341 | 1.000 |  | 0.088 | 0.766 | 1.000 | - | - |  | 12.388 | 0.000 | 0.001 | 0.962 | CCxMMC |
| MMC vs CCxMMC | LOP | 0.612 | 0.434 | 1.000 |  | 0.533 | 0.465 | 1.000 | - | - |  | 26.013 | 0.000 | 0.000 | 0.981 | CCxMMC |
| CC vs MMC | aME | 0.356 | 0.551 | 1.000 |  | 9.391 | 0.002 | 0.007 | 0.951 | MMC |  | - | - | - | - | - |
| CC vs CCxMMC | aME | 1.482 | 0.223 | 0.670 |  | 0.000 | 0.985 | 1.000 | - | - |  | 17.902 | 0.000 | 0.000 | 0.973 | CCxMMC |
| MMC vs CCxMMC | aME | 0.901 | 0.342 | 1.000 |  | 7.484 | 0.006 | 0.019 | 0.939 | MMC |  | - | - | - | - | - |
| CC vs MMC | vLOB | 0.127 | 0.722 | 1.000 |  | 2.187 | 0.139 | 0.417 | - | - |  | 8.869 | 0.003 | 0.009 | 0.948 | CC |
| CC vs CCxMMC | vLOB | 1.484 | 0.223 | 0.669 |  | 0.772 | 0.380 | 1.000 | - | - |  | 12.010 | 0.001 | 0.002 | 0.961 | CCxMMC |
| MMC vs CCxMMC | vLOB | 2.212 | 0.137 | 0.411 |  | 0.251 | 0.617 | 1.000 | - | - |  | 27.985 | 0.000 | 0.000 | 0.983 | CCxMMC |
| CC vs MMC | AL | 0.570 | 0.450 | 1.000 |  | 0.007 | 0.934 | 1.000 | - | - |  | 2.945 | 0.086 | 0.258 | - | - |
| CC vs CCxMMC | AL | 0.726 | 0.394 | 1.000 |  | 0.453 | 0.501 | 1.000 | - | - |  | 12.010 | 0.001 | 0.002 | 0.961 | CCxMMC |
| MMC vs CCxMMC | AL | 0.002 | 0.967 | 1.000 |  | 0.002 | 0.962 | 1.000 | - | - |  | 20.766 | 0.000 | 0.000 | 0.977 | CCxMMC |
| CC vs MMC | AOTU | 2.603 | 0.107 | 0.320 |  | 0.362 | 0.548 | 1.000 | - | - |  | 4.708 | 0.030 | 0.090 | - | - |
| CC vs CCxMMC | AOTU | 1.874 | 0.171 | 0.513 |  | 0.106 | 0.745 | 1.000 | - | - |  | 15.077 | 0.000 | 0.000 | 0.968 | CCxMMC |
| MMC vs CCxMMC | AOTU | 0.228 | 0.633 | 1.000 |  | 1.452 | 0.228 | 0.685 | - | - |  | 33.010 | 0.000 | 0.000 | 0.985 | CCxMMC |
| CC vs MMC | MBCA | 1.398 | 0.237 | 0.711 |  | 0.548 | 0.459 | 1.000 | - | - |  | 4.951 | 0.026 | 0.078 | - | - |
| CC vs CCxMMC | MBCA | 3.413 | 0.065 | 0.194 |  | 2.603 | 0.107 | 0.320 | - | - |  | 9.363 | 0.002 | 0.007 | 0.951 | CCxMMC |

**Table S 30:** Pairwise scaling relationships tests for species differences in neuropil allometry in 10-day old H. cydno, F1 hybrids (CCxMMC), and H. melpomene (n= 34), including slope shifts (β), elevation shifts (α), and major axis shifts relative to rCBR. FDR-corrected p-values, effect sizes (r), and direction of increase (DI) are reported. Values are for tests for grade shifts (α) and shifts along the common axis are only appropriate with a common slope.

32.	R Core Team, *R: A Language and Environment for Statistical Computing* (R Foundation for Statistical Computing, 2025).
